# Minimal social context decouples affective response modalities

**DOI:** 10.64898/2026.04.17.718894

**Authors:** Ruth Judson, Jay Louise Davies, Josie Briscoe, Hélio Clemente Cuve

**Affiliations:** School of Psychology and Neuroscience, University of Bristol

**Keywords:** affect, emotion, social context, coherence, autonomic, facial expressions

## Abstract

Emotions often occur within social interactions where affective cues are partly accessible or inferable by others. This raises questions regarding how and to which degree social context modulates subjective, physiological and behavioural affective responses, as well as their coherence, questions which remain points of tension in emotion research. To investigate this, we measured subjective affective ratings, autonomic sympathetic and parasympathetic activity, and facial behaviour while participants completed an emotion-induction task. In the social-context condition (but not control), participants believed that their video feed was accessible to a potential future interaction partner. Results show that even such “minimal social context” selectively and differentially modulated affective response modalities, characterised by both intensification of autonomic responses and dampening of overt facial and subjective affect. Multivariate dimensionality analysis further identified a cross-modal affective dimension. Notably, social context reduced participants’ coupling with this shared affective response structure, indicating weaker cross-modal coherence. These findings suggest that emotional responding relies on a flexible, rather than rigid, configuration of affective features, likely recruited to meet the socioemotional demands of a given context. This has important implications for understanding the structure and function of emotion, as well as typical and atypical socioemotional responding.

## 1. Introduction

Emotional episodes are thought to coordinate changes across subjective experience, autonomic activity, and behavioural responses often theorised to serve intrapersonal functions. At the same time, these responses are far from private; emotions are often generated and experienced within social interactions or contexts where affective cues such as facial expressions are visible to and directed towards others. Even in the case of autonomic affective cues reflecting internal bodily states, it has been argued that at some level they may also be detectable or inferred by observers (Arslanova et al., 2022; Kret, 2015). Indeed, theoretical disagreements regarding the structure and function of emotion are often emphasised in the literature, there is broad theoretical acknowledgement of the potential for social context in shaping the generation or expression of emotion.

In particular, most emotion theories have attempted to provide an explicit account linking interpersonal social functions and affective responses. Functionalist emotion theories hypothesise that contextual display rules may explain the modulation of otherwise “biologically hardwired” repertoires of affective expression in social contexts (Ekman et al., 1987). Socio-communicative theories of emotion on the other hand, view affective signals as fundamentally directed to influence others (Crivelli & Fridlund, 2018; Fridlund, 1997). Others propose that emotional responses (and related signals) serve a dynamic process of relational alignment inherent to social interaction (Parkinson, 2021). While much of these arguments have focused on the study of facial expressions, social modulation can theoretically extend to the experiential subjective, and embodied features of emotion. Indeed, neither of the aforementioned accounts negates the potential for coordinated and patterned responses across affective modalities. Nonetheless, contemporary critiques of “functional patterning” emphasise that emotions are psychologically and socially constructed ad-hoc and reflect a process of categorisation of necessarily variable patterns of behavioural, subjective and autonomic changes (Barrett et al., 2025). While there are fundamental disagreements between the aforementioned theoretical views, the discussion and adjudication of which is beyond the scope of this work and extensively treated elsewhere (Barrett et al., 2025; Parkinson, 2005; Shiota et al., 2026). Nevertheless, these perspectives converge in expecting social influences on emotional responses. It is therefore surprising that empirical research has largely overlooked how social context influences emotional response patterns across different affective modalities.

From a theoretical standpoint, there are at least three factors that may explain how the structure of emotional responses may be differentially affected by social context. First, response modalities differ in their transmissibility to others, with for example, the overt nature of facial expressions allowing them greater socio-communicative potential than more “private” autonomic and subjective channels. Second, even within the autonomic domain, responses are partly governed by distinct control mechanisms (Berntson et al., 1991; Norman et al., 2014) which may be more or less influenced by social context. Together, these factors may help explain the heterogeneous findings in the literature regarding autonomic and facial expression markers of emotion, which may partly reflect differences in the social orientation of the contexts in which emotional responses are elicited, measured or interpreted (Barrett et al., 2019; Kreibig, 2010; Siegel et al., 2018). The third factor follows directly from this possibility of selective social modulation and concerns whether, and to what extent, subjective, autonomic and behavioural changes are coordinated during emotional episodes, often referred to as the *coherence problem* (Mauss et al., 2005; Van Doren et al., 2021). Any differential social effects on subjective experience, autonomic responses, and behaviour may also alter the degree to which these modalities covary. In this sense, coherence may not be a fixed property of emotional responding, but a context-sensitive feature that tunes affective response and signals to social affordances. However, much of the existing research relies on unimodal and univariate approaches, which are sub-optimal for probing affective coherence. Indeed, multivariate work aiming to decompose patterns of variance underlying affective responses has identified both a coherence factor linking subjective and autonomic responses and additional dimensions unique to physiological activity (Cuve et al. 2023), which may also reflect individual differences (Koppold et al., 2026). Nonetheless, the question of how social context modulates coherence in emotional responses remains unexplored.

Interestingly, evidence of social modulation of socio-cognitive processes largely stems from “non-emotional” contexts. Within social attention research, second-person paradigms have demonstrated that real and implied presence of others alters social behaviour, including how people gaze at others (Cañigueral et al., 2022; Risko et al., 2016). Nonetheless, these findings draw parallels with early work on emotion and facial expression production highlighting “audience effects”, whereby people suppress or exaggerate affective expressions in the presence others (Bruder et al., 2012; Buck et al., 1992). There is, however, a paucity of research examining social context modulation of subjective, autonomic, and behavioural affective modalities collectively. One of the few examples in this direction comes from studies of pain responses, showing that the mere belief of being watched dampened facial expressions, subjective experience and autonomic response (Kleck et al., 1976). However, pain is a unique sensory-affective experience that may not translate to wider emotional responding and social interaction.

The present study therefore explored how social context modulates emotional responding. We used a minimal social context manipulation to induce belief of social observation from a potential unknown interaction partner in half of our participants. We then measured subjective affect (valence, arousal, and emotion category evaluations), autonomic (cardiac and electrodermal activity) and facial behaviour, while all participants completed an emotion induction task. These modalities were selected because they represent commonly used measures of emotional responding across experiential, physiological, and behavioural domains in psychophysiological studies (Azari et al., 2020; Kreibig, 2010; Siegel et al., 2018), while also varying in their degree of overt social transmissibility (Arslanova et al., 2022; Kret, 2015). However, because our aim was to examine social modulation and cross-modal coherence of affective responses, we did not assume any pre-defined prototypical patterns corresponding to a given emotion for subjective, facial behaviour and autonomic responses. This approach is consistent with contemporary evidence of variability in emotion-related facial, physiological, and experiential responses cautioning against one-to-one mappings between emotion and specific response features(Barrett et al. 2019; Siegel et al. 2018; Cuve et al. 2023). Accordingly, unlike most previous studies, subjective emotion categories were defined by participants’ own responses to each stimulus. Similarly, in line with contemporary approaches to dynamic facial signalling, facial behaviour was treated as dynamic patterns of action-unit activity that could vary with both subjective emotion and social context, without presupposing specific AU configurations (Cuve et al. 2026; Barrett et al. 2019) – see Methods.

Building on the aforementioned gaps, we aimed to address two key questions. First, (RQ1) does social context have a differential impact on affective modalities? Given that prior research shows both facilitative and inhibitory effects of social context, we hypothesised that: (H1) social context will differentially modulate affective modalities, operationalised as (non-directional) differences in response patterns between the social and non-social conditions. On the basis that facial behaviour is more overt than autonomic and subjective experience, we also anticipated that: (H2) social modulation would be stronger for facial behaviour compared to subjective and autonomic responses. In light of the mixed evidence for specific emotion profiles in the literature, and differential effects of emotion category on social modulation, we also hypothesised: (H3) a significant modulation of subjective, autonomic and behavioural responses by emotion category and interactions between social context and emotion.

Second, we investigated (RQ2): whether social context modulates the coherence structure of affective responding across modalities. Specifically, rather than testing differences in the magnitude of responses within each modality (RQ1), this question focuses on whether social context alters the degree of covariance across subjective, autonomic and behavioural measures, indexed using a joint multivariate approach (Cuve et al. 2023). In line with H1-3 we also explored whether social context additionally modulated the extent to which these affective modalities covary during emotional induction. This research therefore provides the first systematic examination of the role of (minimal) social context in modulating multimodal affective responses and their coherence.

## 2. Results

We first compared whether social vs non-social groups were matched on baseline characteristics. We found no significant group differences in age (*t*(86.5) = 0.23, *p* = .816; *d* = 0.05), gender distribution (*X*^2^ (2) = 2.39, *p* = .303) or diagnosed mental health conditions (*X*^2^ (1) = 0.02, *p* = .886). Furthermore, there were no baseline differences between alexithymia traits (*t*(82.3) = -1.31, *p* = .192; *d* = -0.29), autistic traits (*t*(80.465) = -0.79, *p* = .433; *d* = -0.18) or self-reported state mood (*t*(84.941) = 0.02, *p* = .988; *d* = 0.00). We also conducted a social manipulation check. A two-sample t-test found that participants in the social observation condition felt significantly more observed than participants in the control condition (*t*(87) = -1.980, *p* = .025; *d* = -0.42). A one-sample t-test further indicated that the mean belief rating of participants in the social observation condition was significantly greater than 6 (higher than the midpoint) on the scale of 0-10 (*t*(42) = 2.01, *p* = .025; *d* = 0.62). This was corroborated by the debrief comments indicating that the social manipulation was acceptably successful. As such, differences between social and non-social condition, cannot be explained by baseline differences between groups. In the following sections, we present the key results of the social modulation of emotion response modalities and coherence.

### 2.1. Effect of social context on individual emotion response modalities

#### 2.1.1. Subjective affect is modulated by emotion, and selectively by social context

Below we present results of social modulation on subjective affective ratings, namely valence, intensity, and categorical emotion experience ratings.

Results of the mixed models predicting valence showed no significant main effect of social context (*F*(1, 102.5) = 0.01, *p* = .936), however there was a significant main effect of subjective emotion (*F*(5, 5439) = 244.96, *p* < .001). Post-hoc comparisons of estimated marginal means (Tukey-adjusted) showed unsurprisingly that happy and neutral emotions were associated with more positive valence than subjective experience of angry, disgust, fear and sad (all *p.adjust* <.0001). There was no significant interaction between social condition and subjective emotion category on valence (*F*(5, 5435.7) = 1.08, *p* = .369). Results are summarised in Fig.2, and full reporting of post-hoc analyses are included in SM3.

**Fig. 1.**
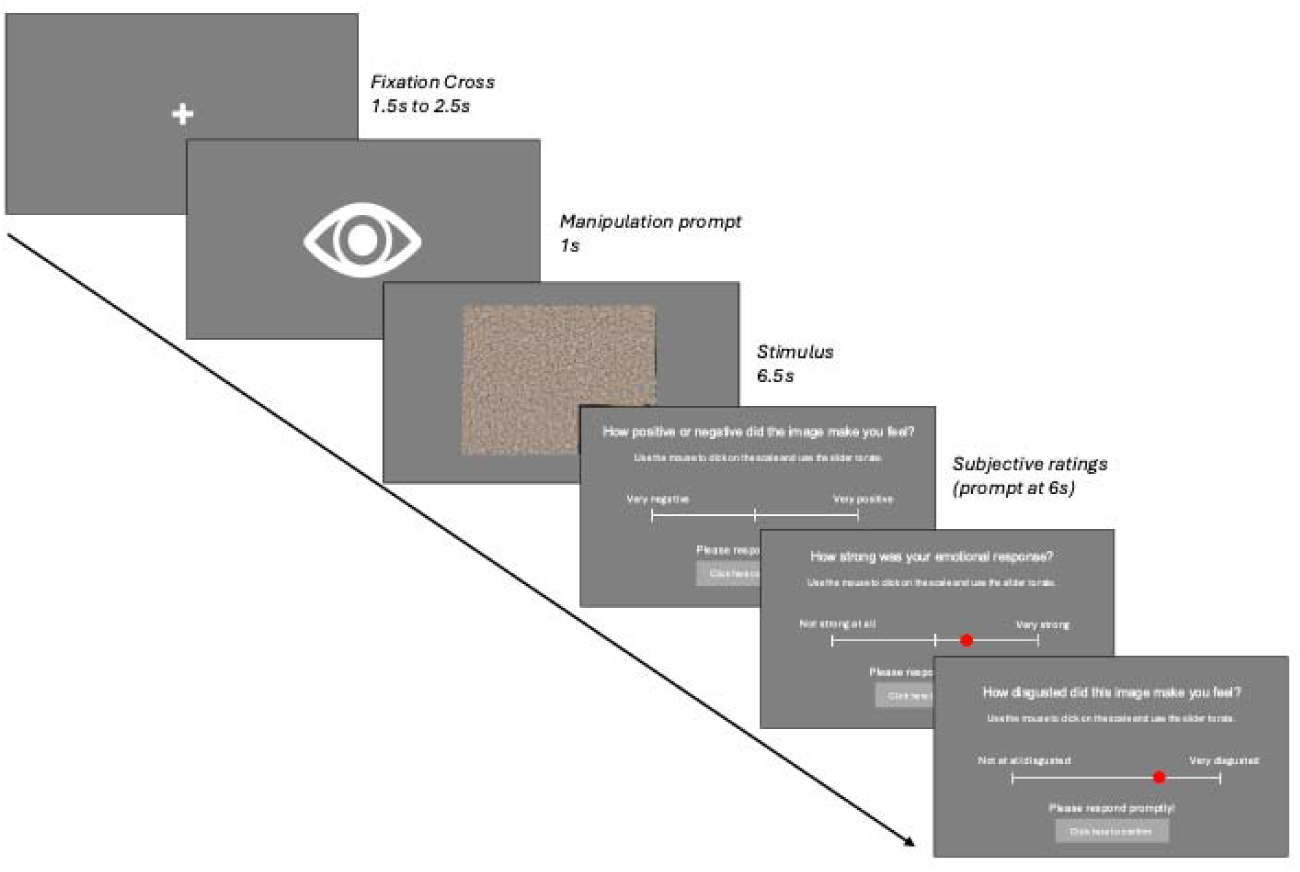
Schematic of Affective Induction and Measurement Task *Note*: An example trial structure. In the social condition only, participants were reminded of the manipulation at the start of each trial after the fixation cross. Autonomic and behavioural recordings were collected throughout the task.

**Fig. 2.**
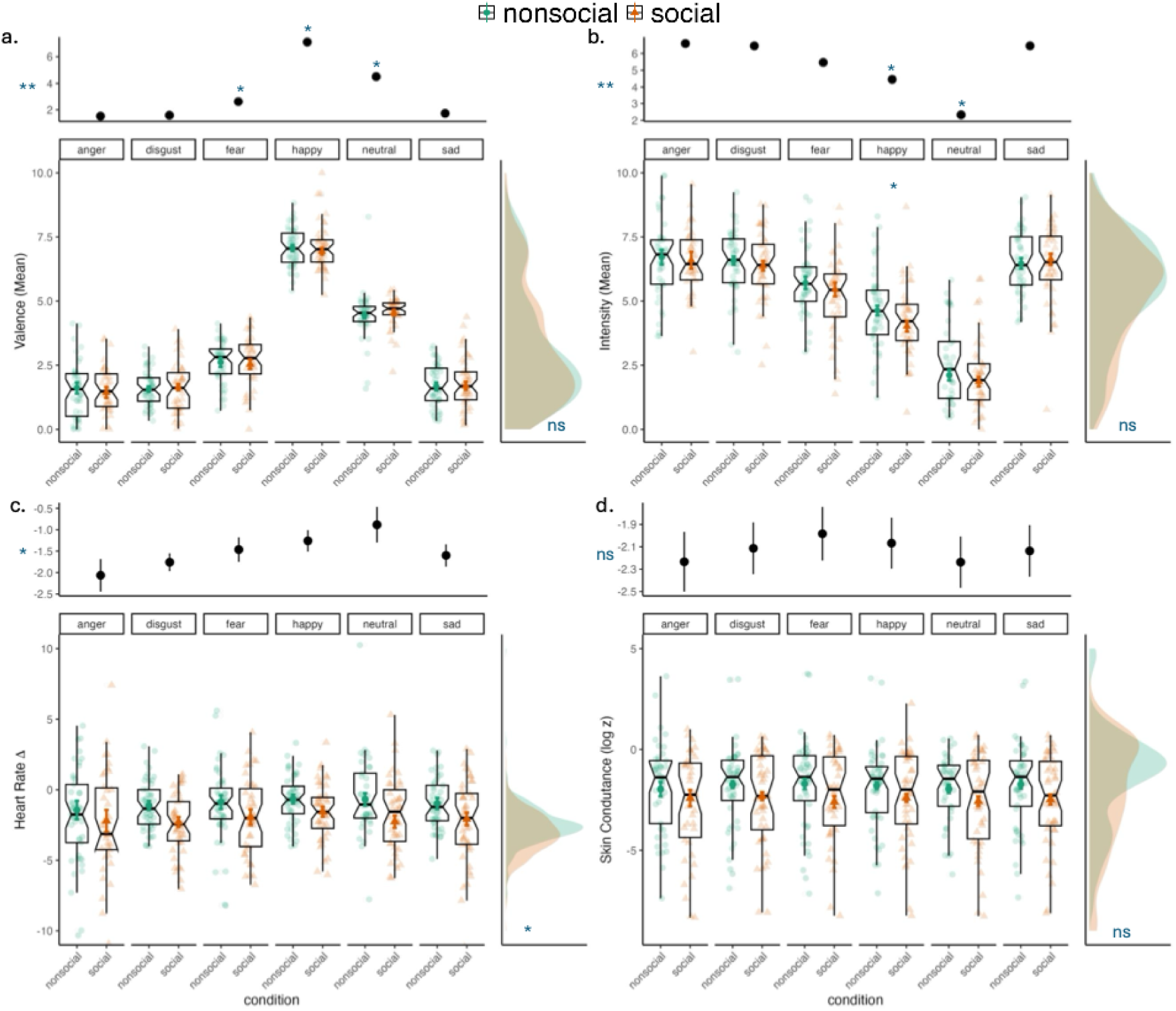
Social modulation of affective modalities Note. Average emotional valence ratings (a), emotional intensity ratings (b), heart-rate change (c), and skin conductance response (d), shown separately for the six subjectively derived emotion categories (anger, disgust, fear, happy, neutral, and sad) and for the non-social and social conditions. The larger central plots display the full emotion category × social context pattern. Participant-level observations are shown as scattered points within each emotion-by-condition cell. Overlaid markers indicate condition means, with error bars representing 95% confidence intervals. The marginal panels provide complementary summaries of the main effects. The top marginal panel collapses across social context and displays marginal means and error bars for each emotion category, corresponding to the main effect of emotion. The right-side marginal panel collapses across emotion category and displays the marginal mean distribution for the non-social and social conditions, corresponding to the main effect of social context. Annotations in the marginal panels indicate the significance of the corresponding main effects; annotations above specific emotion categories indicate involvement in one or more significant pairwise differences. Annotations within the larger central plots indicate specific social versus non-social pairwise comparisons within emotion categories where relevant, such as following a significant social context × emotion interaction. * p < .05, ** p < .01, *** p < .001; ns = non-significant. Axis labels shown on the larger central plots apply to the corresponding marginal plots. Annotations are provided to guide visualisation only; detailed statistics and pairwise comparisons are reported in main text.

Similarly, for intensity ratings, there was no main effect of social context (*F*(1, 94.2) = 1.46, *p* = .23). Like valence, intensity ratings were significantly modulated by felt emotion (*F*(5, 5274.8) = 155.97, *p* < .001). Neutral responses were consistently lower in intensity than all other emotions across both social and non-social contexts, whereas anger, disgust, and sad stimuli were rated as the most intense (all *p.adjust* < .001). Notably, a significant interaction between social context and emotion category did emerge (*F*(5, 5402.5) = 3.71, *p* = .002), where this effect was driven by happy stimuli being rated as more intense in non-social compared to social conditions (β = 0.61, *SE* = 0.2, *p.adjust* = .007).

When looking at felt categorical emotion ratings, social context had no effect on whether participants rated feeling happy (*F*(1, 91.1) = 1.30, *p* = 0.25), sad (*F*(1, 91.2) = 3.18, *p =* .08.), fear (*F*(1, 91.4) = 2.79, *p* = .09), disgust (*F*(1, 91.1) = 1.89, *p* = .17) or anger (*F*(1, 91.4) = 2.25, *p* = .14). Note that for these models, emotion category was not included as a predictor as using the presented emotion to predict corresponding emotion ratings offers little explanatory value (i.e., images from the “happy” category are expected to elicit higher happy ratings by design). Instead, these analyses were interested solely in whether social context modulated participants’ subjective emotional responses. Estimates from each of the emotion slider models were also pooled to allow for a general assessment of the effect of social manipulation on subjective emotional responses (see Methods). Pooled estimates confirmed no effect of social manipulation on categorical emotion experience (β = 0.42, *p* = .12)

In summary, while social context did not significantly influence subjective valence or categorical emotion experience, it partially modulated intensity of happy responses, which were experienced as more intense in non-social contexts, compared to minimal social contexts.

#### 2.1.2. Selective modulation of autonomic reactivity by social context and emotion

The current section presents results investigating social modulation of autonomic affective response, namely HR change (deceleration - acceleration) and SCR.

Fig.2 depicts individual participant and group means for HR changes (BPM) for each emotion across both social and non-social contexts. There was a significant main effect of social condition (*F*(1, 96.8) = 8.49, *p* = .004), such that heart rate deceleration was greater in the social (estimated marginal mean, or EMM = -2.05, SE = .28) compared to the non-social condition (EMM = -.97, SE = .26). There was also a significant main effect of emotion (*F*(5, 714.0) = 2.28, *p* = .045). However, post-hoc comparisons did not reveal significant pairwise differences between subjective emotion categories. Inspection of the 95% CI bars suggests substantial overlap across emotion conditions (Fig.2c). Similarly, the interaction between social context and emotion category was not significant (*F*(5, 5424.9) = 0.78, *p* = .56), suggesting the degree of heart rate deceleration did not depend on the subjective emotion response effect. The social modulation of heart rate deceleration therefore appears relatively uniform across the different emotion responses.

For SCR, there was no significant main effect of social context (*F*(1, 92.3) = 2.27, *p* = .14, see Fig.2d). Similarly, there was no main effect of emotion category (*F*(5, 1258.9) = 0.23, *p* = .95), nor a significant interaction between emotion and social condition (*F*(5, 5322.1) = 1.72, *p* = .13).

In summary, minimal social context selectively modulated autonomic responding, leading to greater HR deceleration, and no significant modulation of SCR.

#### 2.1.3. Selective modulation of facial response components by emotion and social context

Given the complexity of facial behaviour involving changes across multiple AUs and over time, we first identified key underlying spatiotemporal patterns that are diagnostic of facial expression dynamics (Cuve et al. 2026) - see Methods for full details). Specifically, the timeseries for all 17 FAUs were reduced into three interpretable response components representing synergies of AU coordination that differentiates facial behaviours. Component 1 (C1) was characterised by lower face related activity (i.e. associated with AU14 dimpler, AU10 upper lip raise, AU12 lip corner pull and AU06 cheek raise). Component 2 (C2) was characterised by upper face-related activity (associated with AU04 brow lower and AU07 eyelid tighten). Component 3 (C3) was characterised by both lower and upper face related dynamics (associated with AU23 lip tighten, AU09 nose wrinkle, AU45 eye blink, AU20 lip stretch, AU05 eyelid raise, AU02 outer brow raise, AU01 inner brow raise, AU17 chin raise, AU15 lip corner depress, AU25 lips apart and AU26 jaw drop) – see Fig.3. For each component, 38 temporal features were extracted using CMFMTS and subsequently reduced using principal component analysis. The first principal component (PC1) explained the largest proportion of variance (C1: 42.2%; C2: 31.5%; C3: 26.2%) and was therefore retained for subsequent analysis. The component score represents a projection of the original feature set onto the dominant axis of variation and can therefore be understood as reflecting the overall diagnostic dynamic facial pattern. Note that this analysis is agnostic to the emotion, and as such the components do not correspond to specific emotions, but rather to patterns that can be used to differentiate different emotion expressions on the basis of facial dynamics.

**Fig. 3.**
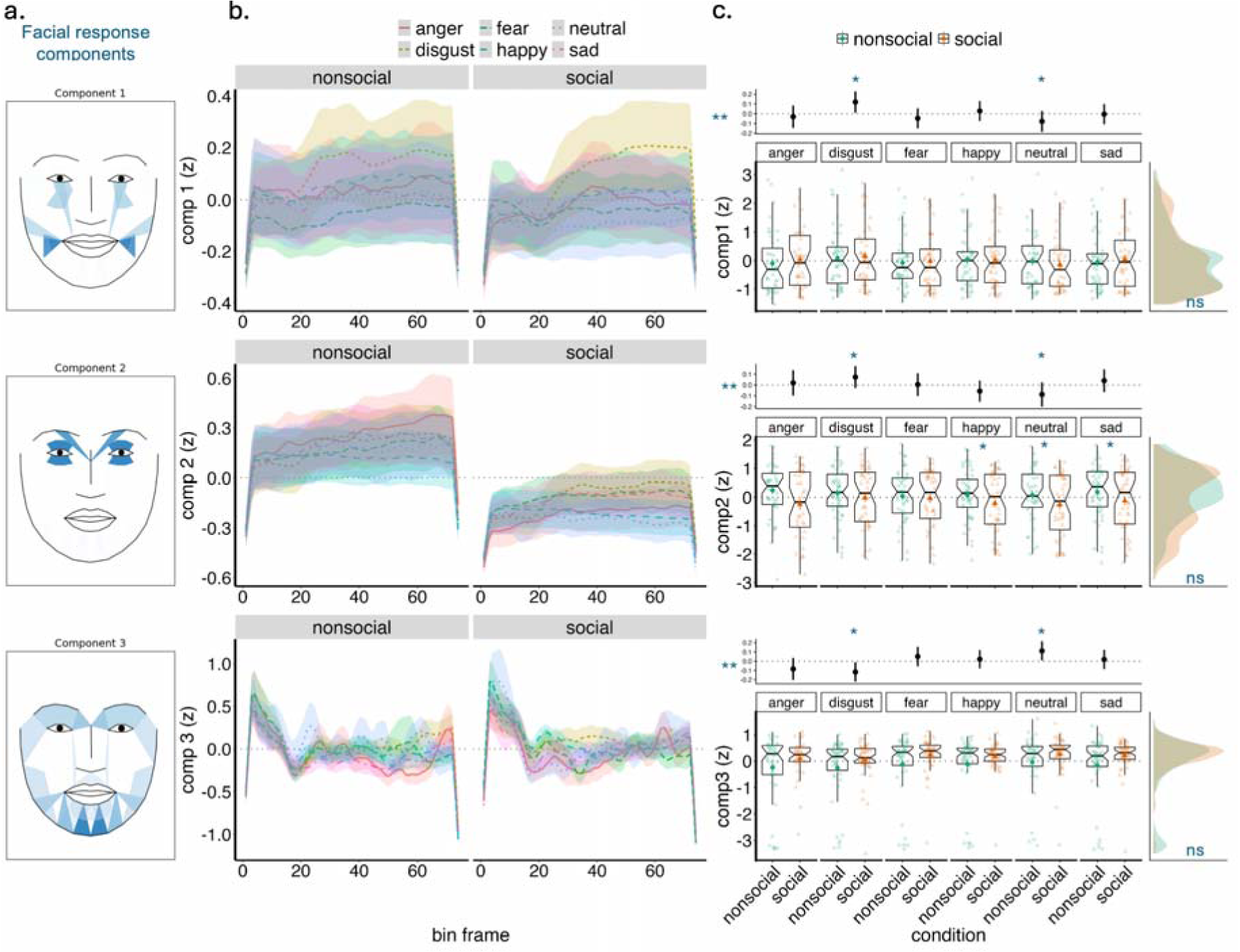
Emotive facial behaviour A. Summary of facial response components. B. Summary of facial response components over time. C. Visualisation of facial response components used for final modelling of social modulation, faceted by the six subjectively derived emotional categories (anger, disgust, fear, happy, neutral, and sad) and non-social and social groups. The larger central plots display the full emotion category × social context pattern. Participant-level observations are shown as scattered points within each emotion-by-condition cell. Overlaid markers indicate condition means, with error bars representing 95% confidence intervals. The marginal panels provide complementary summaries of the main effects. The top marginal panel collapses across social context and displays marginal means and error bars for each emotion category, corresponding to the main effect of emotion. The right-side marginal panel collapses across emotion category and displays the marginal mean distribution for the non-social and social conditions, corresponding to the main effect of social context. Annotations in the marginal panels indicate the significance of the corresponding main effects; annotations above specific emotion categories indicate involvement in one or more significant pairwise differences. Annotations within the larger central plots indicate specific social versus non-social pairwise comparisons within emotion categories where relevant, such as following a significant social context × emotion interaction. * p < .05, ** p < .01, *** p < .001; ns = non-significant. Axis labels shown on the larger central plots apply to the corresponding marginal plots. Annotations are provided to guide visualisation only; detailed statistics and pairwise comparisons are reported in main text. Detailed statistics are provided in-text.

Linear mixed models revealed no significant main effects for social condition across all three components: C1 (*F*(1,92.7) = 0.1, *p* = .75); C2 (*F*(1, 92.7) = 3.35, *p* = .07); and C3 (*F*(1, 93.8) = 2.69, *p* = 0.1). However, there was a significant effect of emotion category for lower face component C1 (*F*(5, 493.8) = 4.27, *p* = <.001), upper face component C2 (*F*(5, 729.3) = 4.41, p <.001), and lower and upper face component C3 (*F*(5, 487.8) = 2.56, *p* = .03). Post-hoc comparisons of estimated marginal means (Tukey-adjusted) confirmed that for all components, the main effect of emotion was largely driven by disgust stimuli. Specifically, lower face AUs (C1) were activated significantly more by disgust stimuli (EMM = 0.33) than all other emotion conditions (fear: EMM = -0.13, β = 0.46, *p* = .01; neutral: EMM = -.12, β = 0.45, *p* = .01; sad: EMM = -0.16, β = 0.49, *p* = .003; and anger: EMM = -0.19, β = - .52, *p* = .04). For C2, upper face AUs were activated significantly more for disgust stimuli (EMM = 0.29) compared to happy (EMM = -.018; β = -.47, p <.001) and neutral (EMM = -.03; β = -.49, *p* = .001) stimuli. The inverse pattern was observed for C3, where disgust stimuli (EMM = -.022) reduced activity for the remaining upper and lower face AUs relative to happy (EMM = 0.12; β = -.33, *p* = .04) and neutral (EMM = 0.19; β = -.4, *p* = .02) stimuli. Additionally, a significant interaction did emerge between social context and emotion for upper face AUS - C2 (*F*(5, 5400.8) = 4.52, p<.001). Tukey adjusted post-hoc comparisons revealed that activity of upper face AUs (AU04 and AU07) was reduced in social contexts for happy (EMM = -.81; β = 1.26, *p* = .02), neutral (EMM = -.78; β = 1.15, *p* = .04) and sad (EMM = -.06; β= 1.169, *p* = .03) emotion responses (for full post-hoc reporting see SM4). In other words, social context reduced expressivity of upper face actions for these emotional responses.

In summary, in relation to RQ1, we find that social context did have differential impacts across the affective modalities in line with H1. Subjective ratings of affect remained largely invariant to our social manipulation, with no differences in valence, intensity or emotion ratings observed between social and non-social contexts. Nonetheless, there was evidence of selective de-intensification of happy responses under social compared to non-social context. In contrast, heart rate measures were significantly impacted by social context, whereby we observed greater heart rate deceleration in social compared to non-social contexts. Facial expressions showed selective sensitivity to social manipulation, with upper face activity reduced in social contexts for low-arousal emotions (happy, neutral, sad), while remaining unchanged for more arousing emotions (anger and disgust).

### 2.2. Reduced cross-modal affective coherence by minimal social context

Beyond analysis of individual response modalities, we also aimed to investigate whether social context modulates the degree of coherence between different affective responses when considered in a common representational affective space (see Methods). A Multiple Factor Analysis (MFA) was conducted on the combined dataset comprising grouped data by subjective ratings, HR changes, and SCR and facial components for all stimuli (see Methods). The MFA solution suggested three primary dimensions accounting for 21.5% of the total covariance across modalities (Dimension 1: 8.4%; Dimension 2: 6.9%, and Dimension 3: 6.2%). Dimension 1 reflected a cross-modal pattern, receiving moderate contributions from all modality groups: heart rate (33.9%), facial dynamics (29.9%), subjective ratings (25.2%), and skin conductance (10.9%) – see Fig.4.

**Fig. 4.**
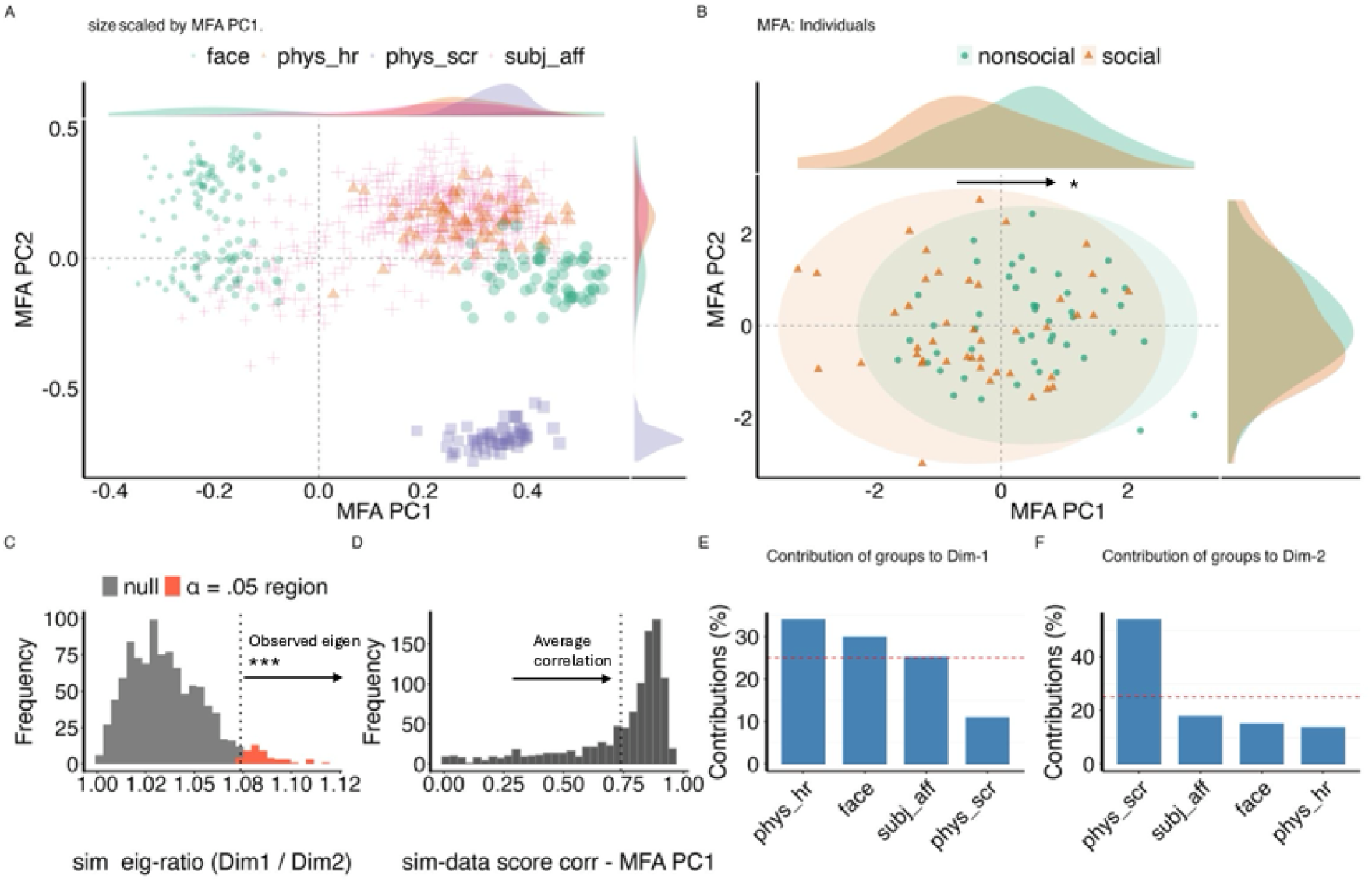
Coherence and affective space A. Plot of affective modalities on a common affective space. Points represent the position of variables for each stimulus across the common dimension. Spread over an axis suggests that variability for that variable is captured by that axis. For example, most variables vary along the x axis, suggesting that this dimension captures cross-modal coherence. B. Plot of individuals by group (social vs non-social) on the common affective space, showing individual differences in the cross-modal affective dimension. C and D Shows validation statistics of the cross-modal dimension based on permutation and bootstrap sensitivity tests respectively. E. Summary of loadings of affective modality and variable groups the first two dimensions. Phys_hr = Heart rate change; Phys_scr = Skin Conductance Response; Subj_aff = subjective affective responses (valence and emotion ratings), face = facial behaviour components.

In contrast, Dimensions 2 and 3 captured more modality-specific covariance structures across participant and trial responses. Dimension 2 was largely represented by skin conductance (53.8%), followed by subjective ratings (17.8%), facial dynamics (14.9%), and heart rate change (13.54%). Dimension 3 was represented by facial dynamics (42.1%), followed by subjective ratings (25.5%), heart rate (20.2%), and skin conductance (12.2%). The relatively balanced contributions across modalities for Dimension 1 indicate coordinated variation across subjective, autonomic, and facial responses, consistent with a cross-modal affective coherence dimension.

To verify that this structure reflected meaningful cross-modal covariance rather than spurious structure, two validation analyses were conducted (see Methods). First, the observed eigenvalue for Dimension 1 exceeded those obtained from permuted datasets simulated with no systematic cross-modal covariance (*p*_(eig_obs> eig_sim_) <.001). Second, bootstrap resampling for stability analysis (see Methods) further demonstrated that the cross-modal dimension was stable. Specifically, the correlation between participant scores and for variable loadings between observed and bootstrapped datasets was on average .73 and .68 respectively, with around 50% of variance shared (see Fig. 4). As such, this dimension was retained as the primary index of cross-modal affective coherence for subsequent analyses examining the influence of social context.

To directly address RQ2, we compared participants’ positions along the cross-modal coherence dimension between the social and non-social groups. Participants in the non-social condition showed significantly higher scores on Dimension 1 than those in the social condition (*t*(84.19) = 3.64, *p* < .001, *d* = . 0.77). This pattern indicates a divergence between conditions, with participants in the non-social context aligning more strongly with the cross-modal coherence dimension (positive scores) than the social group (see Fig.4). This suggests that the presence of minimal social context was associated with reduced cross-modal affective coherence.

We additionally examined whether the quality of the latent structure itself differed between conditions. This analysis addressed the possibility that apparent group differences along Dimension 1 might arise from differences in how well participants’ responses were captured by the shared latent dimension rather than genuine variation in coherence. Specifically, we compared each participant’s contribution to Dimension 1, indexing the degree to which their response variability was explained by the cross-modal structure. Contributions did not differ significantly between conditions (*t*(78.11) = −0.79, *p* = .43, 95%, *d* = -0.17). This indicates that the latent structure was similarly represented across groups and that differences along Dimension 1 reflect differences in the expression of cross-modal affective coherence.

In summary, minimal social context was associated with a reduction in cross-modal affective coherence, reflected in lower alignment with the shared covariance structure linking subjective, autonomic, and facial responses.

Overall, results indicate that even subtle social interactive contexts can produce selective, modality-specific effects rather than global uniform changes in emotional responding. While subjective reports remained largely unchanged, social observation was associated with greater heart rate deceleration, reduced upper-face activity, and reduced cross-modal coherence across affective modalities. Note that several sensitivity analyses addressing potential power issues - by estimating covariance of subjective, autonomic, and behavioural affective responses both within and between individuals as well as assessing structural coherence on a per-participant basis - support the main findings reported here (SM5 and SM9).

Namely, there is a consistent effect of the social versus non-social condition on the cross-modal dimension, alongside notable effects of emotion and interactions consistent with unimodal analysis. Such broad consistency across different variations of the main analysis suggests robustness of the main findings. Crucially, these patterns cannot be explained by differences in visual attention allocation to emotive stimuli (see SM7).

## 3. Discussion

Emotion theories postulate social and communicative functions of affective responses, yet empirical evidence for social modulation of different affective modalities remains limited. Here, we investigated how even a minimal social context modulates individual response modalities, as well as their joint covariation in a common affective space.

### Modality and emotion-specific profiles of social modulation

We observed a nuanced modality-specific pattern of social modulation, indicating both intensification and dampening of affective responses. However, rather than emerging as a consistent main effect of social context across all outcomes, social modulation appeared either as a direct effect of social context within specific modalities (e.g. HR) or as an interaction between social context and emotion (e.g. face and intensity). This pattern is broadly consistent with H1, which predicted differential social modulation across affective modalities. At the same time, social context had no statistically detectable effect on some response features in the present study, including valence ratings and specific categorical emotion ratings or SCR. This suggests that the impact of social context is selective rather than global. Contrary to H2, however, the clearest social-context effect emerged in the autonomic domain, specifically heart rate, rather than in overt facial behaviour. Participants in the social condition showed greater HR deceleration which is theorised to index orienting tendencies to emotionally or salient stimuli (Graham & Clifton, 1966; Lynn, 2013). Although facial behaviour also showed social modulation, this was restricted to upper-face activity for particular emotion categories rather than reflecting a general effect of social context on facial expressivity.

While autonomic modalities did not show robust emotion-specific modulation, the finding that social modulation of facial behaviour varied across experienced emotions, suggests that some emotions may be more socially relevant than others (Fischer & Manstead, 2016; Hess et al., 1995). This is also consistent with social-functional accounts of emotion, which emphasise that expressive behaviour is shaped by interpersonal context and not solely by internal affective state (Fridlund, 1997; Parkinson, 2005). Interestingly, social modulation was tied primarily to upper-face activity which was reduced in social contexts. One possible interpretation is that upper-face cues become particularly salient within social contexts. This would be consistent with findings that observers often preferentially attend to the upper-face region during social perception tasks including emotion perception (Hessels, 2020). From a theoretical perspective considering signaller-receiver dynamics, modulating upper-face cues in “potentially” interactive contexts may therefore reflect the signaller’s sensitivity of the diagnostic salience of these cues to potential observers. Notably, the social modulation of facial responses contrasts recent work showing that social context primarily modulated and amplified lower-face cues with limited changes in upper face cues (Heesen et al., 2024). One plausible explanation is that the co-present, familiar-partner context in Hessens’ study elicited more overt reactions, whereas our minimal implied social context may have elicited a more restrained, self-monitored response. Under uncertain social conditions, individuals may reduce potentially diagnostic signals in order to minimise the risk of mis-signalling or over-committing to an emotional display (Hietanen et al., 2019). Task constraints, such as the use of a chinrest, may also have limited large lower-face movements in the present study. However, because lower-face cues still differentiated subjective emotion categories, as reported in analysis of facial dynamics, this constraint is unlikely to fully account for the limited social modulation of lower-face activity. Moreover, the emergence of modulation of subtle upper-face activity, is consistent with views that that facial responses to emotion are not rigid fixed configurations (Barrett et al., 2019) but are rather flexible cues that can vary with constraints on the face and context. Indeed, it was previously shown that in conditions where lower face is constrained by the need to simultaneously signal emotion and speech content, which reduces the dynamic range for production of clean prototypical facial expressions, signallers accentuate emotion information via the eyes (Cuve et al. 2026). Overall, the observation of expressive emotion modulation even under these minimal, non-interactive and constrained signalling conditions, highlights the pervasive nature of social context effects on affective behaviour.

Interestingly, SCR did not show differentiated responsivity in the social condition relative to the control condition. The divergence between cardiac and electrodermal responses may reflect differences in the underlying control mechanisms of these autonomic systems. Cardiac responses are influenced by both sympathetic and parasympathetic branches of the ANS, whereas SCR primarily reflects sympathetic sudomotor activity (Berntson et al., 1991; Norman et al., 2014). Consistent with this distinction, prior research has reported dissociations between cardiac and electrodermal indices during orienting responses (Barry, 1979, 2009; Latash, 1990). Nonetheless, the observed social difference in HR deceleration still raises the question of why such a pattern might emerge, as well as the potential emotional and functional underpinnings. In line with the idea of social modulation, greater HR orienting to emotional stimuli may reflect increased monitoring and regulation of one’s own responses in the presence of others which is known to activate parasympathetic control (Gross, 2015; Zajonc, 1965). In contrast, SCR (and sympathetic) activity may be more susceptible to dampening in social contexts (Boucsein, 2012; Bradley et al., 2012; Norman et al., 2014).

### Cross-modal coherence, social effects, and affective responses

The observed cross-modal coherence dimension provides partial support for the view that affective responses are partly mobilised as an integrated emotive episode. However, this coherence was limited in magnitude (21.5% variance explained), indicating substantial modality-specific structure alongside shared variance (Cuve et al. 2023). Combined with the observed differences in coherence between social conditions, these findings suggest that variability in autonomic and behavioural correlates of emotion is meaningful rather than noise. In particular, emotional responses may involve dynamic reconfiguration of affective modalities in order to meet the interpersonal demands associated with a given situation (Fischer & Manstead, 2016; Quigley & Barrett, 2014). For example, fear may require a rapid, high-arousal, action-oriented configuration when escaping a physical threat. However, in a social-interactive context it may instead be beneficial to dampen overt behaviour in order to avoid signalling vulnerability while maintaining physiological readiness. By the same token, neutral or ambiguous responses may be maintained more rigidly in social settings to minimise the risk of inadvertently signalling unintended affective states during interaction.

Taken together, the modality- and emotion-specific social modulation of affective responses and coherence calls for an explicit consideration of the social and coherence-related functions of affective responding. Although speculative on the basis of the present study, these findings naturally raise questions about mechanisms underlying atypical affective responding and social interaction, which often co-occur. Are these difficulties better understood in terms of differences in the social modulatory functions of affective responses and cross-modal coherence? Furthermore, if looser coupling between affective modalities might index a flexible response mode (e.g. differential responding to social contexts), might differences in affective responding and coherence in atypical conditions reflect functional compensatory patterns rather than generalised “deficits”?

Despite these insights, there are several important limitations to acknowledge in the current study. First, the social manipulation was minimal and non-interactive, which may have constrained the nature of social-affective modulation. It is unclear how well these patterns generalise to stronger or reciprocal interaction contexts. Second, the study did not directly account for potential regulatory mechanisms involved in affective responding and how they may interact with different modalities. Third, differences in the affective signal-to-noise ratio across modalities may influence the patterning of emotional responses and their apparent coherence, potentially contributing to modality-specific effects. Fourth, although the present study included multiple autonomic and behavioural indices, future work could extend this approach by incorporating additional physiological measures such as pupil size to further characterise cross-modal affect coherence (e.g. respiration, pupil).

### Conclusion

Social context shapes affective responding in a modality- and emotion-specific manner. Social context is associated with lower cross-modal coherence across experiential, physiological, and behavioural responses to emotion induction, suggesting flexible rather than rigid emotion features may be recruited to deal with varying socio-emotional demands. These findings highlight the need to consider the theoretical implications and functional significance of socially modulated and flexibly coupled affective systems for understanding the basis of emotion and social response.

## 4. Method

### 4.1. Participants

98 participants recruited from the University of Bristol and surrounding community took part in the study for course credit or small voucher reward. Following exclusions (see Measures and data processing) the final sample consisted of 90 participants [Mean age = 21.54 (18-51); 70 (76.1%) female, 21 male (22.8%), 1 non-binary (1%)]. Given that determining sample size for multimodal emotion studies is challenging due to the prevalence of underpowered studies (Kreibig, 2010; Siegel et al., 2018), the current sample size was informed through a combination of estimates from previous work (Cuve et al. 2023) and constraints associated with recruiting participants for a multimodal study of this nature. To further evaluate the adequacy of the achieved sample size, we conducted a simulation-based sensitivity analysis of the primary social context contrast (see SM10). This analysis suggested that the final sample was likely sufficient to detect small social context effects in the primary unimodal analyses, although these estimates should be treated as approximate given the complexity of variance components, correlation structures and data-driven factors that could not be specified a priori. Nevertheless, the resulting sample is relatively large compared with much of the published literature. The study was conducted in accordance with ethical principles for research involving human participants (the revised 2013 Declaration of Helsinki) and was approved by the University of Bristol Research Committee (approval code: 15644).

### 4.2. Procedure and task

Upon arriving to the lab, participants completed the informed consent process and were set up for autonomic, eye-tracking and facial behaviour recordings. Next, a two-minute rest period was used to ensure that bodily signals were at baseline before completing the affective response task in which subjective, autonomic and behavioural responses were measured. This was followed by a series of demographic and psychometric questionnaires, after which participants were debriefed. The entire task took approximately 1.5 hours and was completed in a light and temperature controlled soundproofed room.

#### 4.2.1 Affective response task

The task consisted of 60 trials presented in a random order for each participant across two blocks (30 trials each). Stimuli were selected from standardised datasets of emotion-eliciting images, mostly from the International Affective Picture System (IAPS; n = 44; (Lang & Bradley, 2007) and from disgust specific databases which are not well represented in IAPS (n = 10 from DIsgust-RelaTed-Images database, DIRTI; (Haberkamp et al., 2017). A list of stimuli IDs and descriptions from the IAPS and DIRTI databases are provided in the Supplementary material (SM1). The normative valence, arousal and categorical emotion strength scores for IAPS images from previous studies (Libkuman et al., 2007) were used to ensure that images were distributed equally across emotion conditions (happy, sad, fear, disgust, anger and neutral). Four additional images were selected as practice stimuli and to determine acceptability of the level of emotion induction to participants. Each trial began with a fixation cross (1,500ms - 2,500ms), followed by the stimulus (6,500ms), and subjective affective ratings (see Fig.1). The experiment was presented in PsychoPy version 2022.2.5 (Peirce et al., 2019) running on an Intel Core i7 Windows 10 computer with 32GB RAM. Stimuli were 1024 x 760 pixels and were displayed on a 24” (1920 x 1080 pixels) monitor, thus measuring approximately 24 x 18 degrees of visual angle at a viewing distance of 65cm, at a 60 Hz fixed refresh rate.

#### 4.2.2. Social observation manipulation

In order to investigate the effect of social context on emotional responding, while preventing carryover effects, participants were pseudo-randomly assigned to one of two conditions: a “social” condition in which task instructions aimed to induce a belief of being observed by another participant, or a non-social observation (or control condition), in which there was no explicit instruction that they were being observed by another participant. Participants in the social observation condition were informed that the monitor’s webcam would be recording them throughout the task, and that a participant in a parallel study (who they might interact with at a later stage during the study) would watch their video feed and answer some questions. To ensure plausibility of the manipulation, the participants own camera feed was left visible to them at the start of the experiment while the experimenter was conducting the initial setup. All participants in the social context condition were verbally presented the social observation manipulation at the beginning of the session, and again via screen instructions at the start of each block of 30 trials. Note that participants did not believe that their subjective ratings would also be visible to an observer. The social manipulation was deliberately kept simple with minimal contextual detail to avoid unintended biases that might arise relating to potential interactants. Similar minimal social context manipulations have been shown to reliably influence behaviour in previous research (Cañigueral et al., 2022; Risko et al., 2016). Moreover, using a minimal context provided a stringent test of social modulation, as observing significant modulation under such subtle conditions would be suggestive of a more fundamental role of social context effects on affective response and coherence. To prevent the salience of social manipulation becoming dormant (Nasiopoulos et al., 2015), a schematic image depicting social observation was presented for 1,000ms at the start of each trial (Fig.1). Note that all participants had consented to video recording and eye-tracking in both conditions. Consequently, participants in the control condition were informed that this equipment would be recording to check their distance to the screen to ensure correct stimuli presentation. As a sanity check, at the end of the task all participants were asked ‘to what extent did you feel observed throughout the task?’. Participants in the social condition were further asked ‘to what extent did you believe that you were being observed by another participant via a video livestream?’. Both questions ranged from 0 (‘not at all observed’/’didn’t believe it at all’) to 10 (‘very observed’/’entirely believed it’).

### 4.3. Measures and data processing

#### 4.3.1. Subjective affective ratings

For each trial, participants responded to a series of questions about their subjective affective experience on a sliding scale. These included valence (‘how positive or negative did the image make you feel?’; 0 = ‘very negative’; 10 = ‘very positive’); and intensity (‘how strong was your emotional response?’, 0 = ‘not strong at all’, 10 = ‘very strong’). They also rated categorical emotional response intensity experienced for 5 labels: happy, sad, fear, disgust and anger (e.g. ‘how [happy] did the image make you feel?’, 0 = ‘not at all [happy]’,10 = ‘very [happy]’). All seven variables together form a variable group henceforth referred to as subjective affective experience.

To categorise emotion for latter analyses, we decided that it would be more appropriate to capture the participant experience rather than using the normative emotion ratings provided with the stimulus sets (see SM1 and SM2). For each trial, the highest score across the five discrete emotion sliders for each participant and trial were used to class their subjective dominant emotion category. A trial was classed as ‘neutral’ if the maximum index score was below 4.5; this was chosen based on exploratory visualisations of frequency distributions of the maximum indices. Visualisation of subjective emotion categorisation shows notable participant by participant variation in the distribution of emotion categories (See SM2, Fig.S1). Nonetheless, there were no differences in emotion categorisation between the social observation and control conditions (see section 3.1.1).

#### 4.3.2. Autonomic measures

Galvanic skin response (GSR) and cardiac activity were recorded throughout the task using Shimmer Sensing units controlled via ConsensysPro software. Specifically, GSR was recorded using the Shimmer GSR+ unit sampling at 128 Hz. The data were collected via reusable velcro-strap electrodes attached to the index and middle fingers of the non-mouse hand: fingers were first cleaned with water before electrodes were attached. Cardiac activity was recorded using a Shimmer ECG unit sampling at 250 Hz, collected bilaterally via disposable surface electrodes attached on the right collarbone and left hip.

GSR data were pre-processed using a Continuous Decomposition Analysis (CDA) Ledalab custom script in MATLAB (Benedek & Kaernbach, 2010). An analysis window of 500-7000ms after the stimulus onset was chosen for analysis so that the entire response from onset to peak was captured in full. The adaptive smoothing filter and default criteria were applied in Ledalab. The time integral of the phasic response was used as the GSR variable, as it represents SCR more accurately. SCR was log-transformed before analysis to adjust for skewness.

Electrocardiogram (ECG) data were pre-processed using the PhysioData Toolbox (SjakShie, 2023) in MATLAB. Raw ECG signals were filtered using a 1 Hz high-pass and 50 Hz low-pass filter, after which R-peaks were automatically detected. Misidentified or missing peaks were visually inspected and manually corrected when their locations could be reliably determined from waveform morphology and inter-beat interval (IBI) patterns. Only 0.46% of accepted R-peaks were manually defined, with the remainder detected automatically using the toolbox’s default algorithm. Mean heart rate (HR; beats per minute) and inter-beat interval (IBI; seconds) were calculated for pre-stimulus and stimulus epochs (7s each). Baseline-corrected change scores were then computed by subtracting pre-stimulus values from stimulus values to index stimulus-evoked cardiac responses. Higher (positive) HR change scores and lower (negative) IBI change scores indicate cardiac acceleration, whereas lower (negative) HR change scores and higher (positive) IBI change scores indicate cardiac deceleration. Changes in HR and IBI were selected as outcome measures because they are widely used indices of autonomic activity in emotion induction research (Azari et al., 2020; Kreibig, 2010; Siegel et al., 2018)

#### 4.3.3. Facial responses

A video of participants’ faces was captured using a webcam integrated with the screen monitor, sampling at 30hz. Video recordings were recorded for every trial and processed offline using OpenFace (Baltrusaitis et al., 2016) for automated facial action unit detection (Ekman & Friesen, 1978). OpenFace provided timeseries estimates for the presence and intensity of seventeen AUs covering a range of muscle movements in the eye, lip, chin, cheek, eyebrow and nose regions. Based on exploratory visualisations, AU intensity patterns contained some jitter likely indicative of small degree of noise inherent to automated facial tracking. Therefore, all AU intensities were smoothed using centred 4-window moving averages with zero-padding to preserve the underlying trend without losing dynamic changes. Because the aim was to characterise how facial behaviour is flexibly modulated by emotion and social context, rather than to test whether participants produced prototypical facial expressions, we did not impose predefined AU-emotion mappings.(Cuve et al. 2026; Barrett et al. 2019). Furthermore, unlike the subjective and autonomic measures, where there is single or small number of derived variables that are well characterised, facial behaviour has a less well-defined response profile and includes changes in several action units over time. Therefore, we used a data-driven spatiotemporal approach to identify latent patterns of co-occurring facial movements and their temporal dynamics (Cuve et al. 2026). This involved three key steps: First, a dimensionality reduction approach was used to identify the key spatiotemporal patterns from the timeseries of facial AUs. Importantly, the aim of this analysis was not to recover canonical facial configurations associated with discrete emotions, but rather to identify latent patterns of co-occurring facial movements that may be flexibly recruited across emotional and social contexts. These AU patterns are visualised using facial heatmaps by feeding basic AU values from the dimensionality reduction into shape-normalised landmarks (Cheong et al., 2023). Second, timeseries features for each spatiotemporal component were extracted, to characterise the diagnostic cues embedded in the dynamics of facial patterns. This was done using CMFMTS method which captures theoretically interpretable timeseries features, such as complexity, temporal autocorrelation and curvature (Baldán & Benítez, 2023). Third, to obtain a parsimonious summary measure representing the latent facial behaviour dynamics, principal component analyses (PCA) were applied on each component’s groups of features. The resulting component score was then used for the linear mixed models (see Statistical analyses and modelling).

Given the minimal and non-interactive context and experimental setting, facial expressivity was notably subtle. However, participant-level sensitivity checks of AU timeseries data confirmed that the facial data included measurable facial activity and temporal variation across participants. Complementary sensitivity analyses using predefined AU-emotion groupings are reported in SM8 and Table S14-16; these analyses were based on average AU activity and did not include the spatiotemporal feature-extraction and dimensionality-reduction steps used in the primary facial analysis.

#### 4.3.4. Eye-tracking

Given that visual attention allocation to emotion inducing stimuli may influence affective responding, we captured gaze behaviour on each trial. Eye- movements were measured using a Gazepoint GP3-HD eye-tracker sampling at 150Hz (Cuve et al. 2022). Participants were positioned 60-65cm away from the screen/eye tracker which had a visual angle of 30°, and a chinrest was used to minimise excessive head movements. A 9-point calibration and validation procedure was used and repeated until the mean offset (error) between these gaze positions was below 100 pixels (2.4 degrees of visual angle). Eye-gaze data were pre-processed using custom scripts to clean and remove invalid gaze samples (e.g. falling outside screen coordinates or flagged as invalid by the eyetracker; (Cuve et al. 2022). Gaze metrics were derived using GazePath (van Renswoude et al., 2018) which applies a data-driven adaptive threshold to classify fixations and saccades, accounting for individual and trial-by-trial differences in gaze quality. Gaze data were used simply for sensitivity checks to assess whether social and non-social groups differed in their visual processing of emotion-inducing stimuli. Note that although pupil size was technically available from the eye-tracking recordings and can provide an additional index of autonomic responding the present study was not designed or optimised for pupillometry. In particular, the complex affective images varied in low-level visual properties which strongly influence pupil responses. This is especially problematic because affective and psychological modulations of pupil response typically account for only a small fraction of overall pupil-size variance, relative to visually driven changes (Mathôt, 2018).

#### 4.3.5. Psychometric questionnaires

A number of questionnaires were used to assess and control for between subject differences in emotion-relevant traits known to impact affective responses and social interaction. This included alexithymia and autistic traits (Bird and Cook 2013; Cuve et al. 2021), as well as depression, stress, and anxiety (Sloan et al., 2017). Alexithymia was measured using the Toronto Alexithymia Scale, measuring difficulty identifying feelings, describing feelings, and externally oriented thinking (Bagby et al., 1994). Autistic traits were quantified using the abridged version of the Autism Spectrum Quotient (AQ-28; (Hoekstra et al., 2011). Additionally, the Depression, Stress and Anxiety Scale (DASS21) was also used (Lovibond & Lovibond, 1995). All questionnaires were scored according to the original scoring procedure and showed good internal consistency in the current sample (Cronbach alpha >.8).

#### 4.3.6. Data exclusions and imputation

To maximise statistical power for individual and joint analyses of affective modalities, data exclusion was applied on a per-signal/variable group basis, ensuring that individual trials met predefined quality thresholds and that participants contributed at least 80% valid data for a given signal. For each signal, trials were excluded if SCR values exceeded ±3 standard deviations from the participant’s mean phasic response across trials, if HR fell outside a physiologically plausible range (30-200 BPM) or exceeded ±3 standard deviations from the participant’s mean HR across trials, and if facial movement tracking success fell below 80%. Note that subjective affective data were always complete. Overall, this resulted in the exclusion of only ∼5% of trials. Eight participants were excluded entirely from facial analyses due to missing most or all face data as a result of technical issues with the camera recording. Overall, the final sample for joint analysis contained 90 participants, with 43 (48.3%) in the social observation condition and 47 (51.7%) in the control condition. However, the number of participants and trials varied slightly across analyses of specific signals. For the joint multivariate analysis, participant data for all modalities were further excluded if any given signal (based on the specific criterion above) had been removed. Allowable missing or invalid trial data were then imputed to create a complete data matrix for coherence analysis. Joint data imputation was performed using the missMDA package in R using a multiple factor analysis (MFA) method which accounts for the grouped structure of the affective data, i.e. subjective, autonomic, and facial behaviour respectively (Husson et al., 2018; Josse & Husson, 2016). To verify that the imputation procedure did not distort the data structure, the imputed dataset was compared with the original dataset containing missing values using paired-samples t-tests of variable means. These comparisons were conducted solely as a descriptive check of the imputation procedure and were not part of the primary inferential analyses. No differences were observed in the mean values of any variables between the pre- and post-imputation datasets (all p > .95).

### 4.4. Statistical analyses and modelling

#### 4.4.1. Linear mixed model analysis: the effect of social modulation on each affective modality

To address RQ1, a series of linear mixed-effects models (LMMs) were conducted to evaluate how social context (social vs. non-social) influenced emotional responses across modalities. Models were performed separately for each affective modality: subjective ratings (valence, intensity, and emotion ratings), autonomic responses (SCR, HR), and facial behaviour. Apart from emotion rating models, models included social context, emotion category, and their interaction as fixed effects. Participant and stimulus were included as random effects to account for participant and stimulus-related variability. The models had the general form:

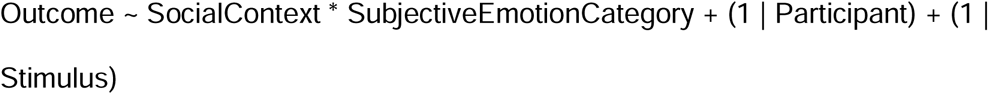

In this format, the asterisk (*) denotes both main fixed effects of social context and emotion, as well as the interaction between them as per lme4 notation. The random-effects structure was determined a priori, prioritising theoretically justified structures while avoiding overly complex specifications (Bates et al., 2015; Meteyard & Davies, 2020). Note that we additionally evaluated models including by-participant random slopes for emotion category:

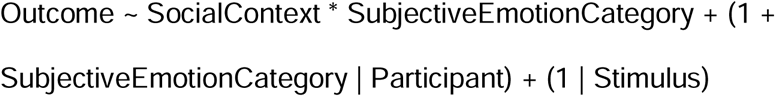

These models produced convergence warnings, singular fits and/or boundary estimates indicating that additional complexity was not reliably supported by the data. Importantly, the substantive pattern of findings was unchanged. We therefore report results of the random-intercept models, with comparisons between model specifications provided in the Supplementary Materials (SM8, Table S13).

All models were fit using maximum likelihood estimation and were implemented using the lme4 package in R (Bates et al., 2014) with inferential tests obtained using lmerTest (Kuznetsova et al., 2017). Statistical significance was evaluated using Type III F-tests with Satterthwaite approximations, and estimated marginal means and pairwise comparisons were calculated using the emmeans package (Lenth et al., 2018). For emotion rating models, the same general model structure was used but subjective emotion category was excluded as a predictor to avoid circularity, as these ratings were derived directly from the subjective emotion slider measures themselves. Additionally, β coefficients for the social versus non-social contrast were extracted from each model to create a pooled social effect across emotions.

Because effects could occur in opposite directions across different emotion ratings, the coefficients were squared prior to pooling then averaged, according to the below formula.

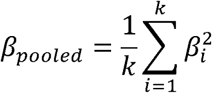

Standard errors were pooled using the same procedure described above, and a z statistic was calculated as the ratio of the pooled estimate to the pooled standard error according to the formula below for inference testing.

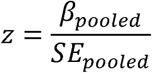

The corresponding two-tailed p-value was derived from the standard normal distribution providing an overall test of whether social context significantly influenced subjective emotional responses across the set of emotion rating models. Note that, where the degrees of freedom contain decimal points, it indicates the use of approximated degrees of freedom, a correction for unequal variance, or estimated degrees of freedom for multilevel models as noted above.

#### 4.4.2. Cross-modal affective coherence analysis: multiple factor analysis

To address *RQ2*, how social context modulates the degree of coherence between affective modalities, we implemented a Multiple Factor Analysis (MFA) to identify latent dimensions capturing shared variance across modalities (Cuve et al. 2023). MFA was chosen for this purpose because it allows the joint analysis of predefined groups of variables while balancing their contributions to the global solution. This is important in the present study, as subjective ratings, autonomic responses, and facial behaviour constitute distinct affective modalities with different numbers of variables. The MFA therefore provided a common representational affective space from which differences between social and non-social experimental conditions could be quantified. Note that instead of using individual AUs for the face behaviour, we used the components scores summarising key spatiotemporal patterns in facial response derived earlier (see 2.3.3. facial responses). The dataset was organised such that each row corresponded to one participant (N = 90), while columns represented responses to each stimulus across modalities (60 stimuli × 4 modalities). Variables were grouped into four modality blocks for the MFA: subjective ratings (7 variables per stimulus), facial components (3), heart rate (1), and skin conductance (1). Across the 60 stimuli, this resulted in a total of 720 variables, yielding a data matrix of 90 participants × 720 responses. The MFA was implemented using the FactoMineR package and applied a singular value decomposition (SVD) of the weighted concatenated data matrix following block normalisation. This is useful in our case as this decomposition can be performed in the observation space, meaning the procedure remains well defined even when the number of variables exceeds the number of observations (Abdi et al., 2013). Nevertheless, to address the potential concern regarding the high dimensionality of the data matrix for the sample size, we conducted two reduced-dimensionality sensitivity analyses in which the data were either reshaped (see SM5) or aggregated (see SM9) so that the number of observations exceeded the number of features. Both sensitivity analyses produced findings convergent with the main analysis reported in the paper.

##### 4.4.2.1. Operationalising and validating cross-modal coherence

To operationalise coherence, we defined the first MFA dimension as the cross-modal coherence axis capturing the largest proportion of variance if it received substantive contributions from all modality groups relative to other dimensions. Under this interpretation, Dimension 1 reflects coordinated variation across subjective, autonomic and facial responses to emotion induction. As such, higher values in this dimension can be interpreted as expressing a stronger degree of coherence (or affective modality covariation). Before testing whether coherence differed between social and non-social contexts, we conducted two validation analyses to ensure that Dimension 1 captured meaningful cross-modal structure rather than spurious latent structure. First, we performed a permutation test under the assumption that the coherence dimension reflects genuine structure, and as such it should be rare to observe a dimension explaining a similar or greater proportion of variance when the relationships between variables are disrupted (e.g. in a null distribution). We generated 1,000 permuted datasets by randomly shuffling values within each variable column, thereby preserving the marginal distribution of each variable while breaking correlations between modalities. Second, we conducted a bootstrap analysis (1,000 iterations) to assess the stability of the cross-modal coherence dimension. Stability was assessed at two levels: variable loadings and participant scores. For each bootstrap iteration, we resampled participants with replacement and recomputed the MFA. To ensure consistency across iterations, we aligned the sign of the bootstrapped dimension to the original observed solution. To assess loading stability, we correlated the variable coordinates on Dimension 1 from each bootstrap iteration with the observed coordinates. To assess participant score stability, we used the variable weights from each bootstrapped model to calculate scores for the entire original sample; we then correlated these predicted scores with the original observed scores. Stability was quantified using the mean correlation across all iterations, with higher values (closer to 1) indicative of higher coherence.

##### 4.4.2.2. Comparing coherence by social condition

To directly examine whether social context modulated coherence, we tested whether individuals’ positions along the coherence dimension differed between the social and non-social context groups. Because differences may also arise from the degree to which participants’ response patterns are meaningfully reflected by the latent coherence structure, we additionally examined whether the quality of dimensional structure differed between conditions operationalised as the proportion of variance explained by Dimension 1, by back-projecting participants data onto the learned affective space. Group differences in both the participant scores on Dimension 1 and the degree to which their responses are well explained by Dimension 1 were first evaluated using independent-samples t-tests. A sensitivity permutation test was also conducted where social context labels were randomly shuffled across participants (5,000 iterations), and the test statistics (difference between social vs non-social labels) were recomputed for each permutation to generate a null distribution against which the observed group differences were compared to derive a probability value.

#### 4.4.3. Pre-registration, deviations, and transparency

The study analyses were preregistered (AsPredicted #188,683: https://aspredicted.org/ndr3-pty6.pdf). A number of motivated deviations were made for methodological accuracy and theoretical considerations which we note for transparency. The final analyses reported here retained the same core aims and hypotheses (i.e. differential modulation of affective modalities by social context). First, gaze was omitted from the analyses of affective behavioural responses because it primarily indexes attentional allocation rather than affective responding per se, allowing closer conceptual alignment with the study’s focus on affective signals and wider literature (Siegel et al. 2018; Cuve et al. 2023). Second, the preregistered plan to incorporate MFA-derived components capturing coherence and divergence within mixed-effects models was slightly revised. The originally proposed approach was not well suited to the hierarchical structure of the data, which included both participant- and trial-level variation, which violates MFA assumptions of independent observations. Modelling this dependency structure within the preregistered framework was not straightforward and remains an active area of research (Abdi et al., 2013). Instead, we implemented MFA using a wide-format structure in which trials were represented as separate columns grouped by affective modality for the key variables. This approach preserved MFA’s ability to capture cross-modal covariance while better respecting the assumptions of the model. Importantly, the originally preregistered analysis was also conducted and yielded convergent results (see SM5, Table S8), supporting the robustness of the findings. However, we reasoned that given the risk of confounding the within and between subject variance in the pre-registered approach, which may inflate Type I and Type II errors, the reported analysis provides a more accurate estimate for any effect of social vs non-social condition. Additional analyses and sensitivity checks were conducted a posteriori and are noted as such; these are used to support the primary analyses rather than serving as a central focus in their own right. Code and data will be available on GitHub https://github.com/hcuve/social_emophys_paper and OSF respectively https://osf.io/cpkam/overview?view_only=16a245801d0a47a481061c11cde7318a.

## Declarations

Competing interests: None

## Data availability

Raw and processed anonymised data will be made fully public with publication on OSF: https://osf.io/cpkam/overview?view_only=16a245801d0a47a481061c11cde7318a

## Code availability

Code for processing and analysis will be made fully public with publication on GitHub https://github.com/hcuve/social_emophys_paper

## Author contributions

Ruth Judson: Conceptualization, Formal analysis, Investigation, Data Curation, Writing - Original Draft, Formal analysis, Project administration, Validation, Visualization Jay Louise Davies: Formal analysis, Methodology, Writing - Review & Editing, Validation, Visualization Josie Briscoe: Conceptualization, Supervision, Writing - Review & Editing, Validation, Visualization Hélio Clemente Cuve: Conceptualization, Methodology, Software, Validation, Formal analysis, Investigation, Resources, Data Curation, Writing - Original Draft, Writing - Review & Editing, Visualization, Supervision, Project administration, Funding acquisition

## Ethics statement

The study received ethical approval from the University of Bristol Research Committee (approval code: 15644).

## Informed consent

All participants gave written informed consent in accordance with ethical principles for research involving human participants (the revised 2013 Declaration of Helsinki) and the University Research committee.

## Funding Statement

This project was partially funded by a Wellcome Trust Award 311793/Z/24/Z

## Notes

### Competing Interest Statement

The authors have declared no competing interest.

### Summary of Updates

fixed typos, provided additional clarifications, improved quality and annotation of figures

