## Supplementary material for "Minimal social context decouples affective response modalities"

### SM1. List of Stimuli

*IAPS images:* Selected from the International Affective Picture System (Lang & Bradley, 2007), with normative emotion categories provided by (Libkuman et al., 2007) as well as responses from our previous study (Cuve et al., 2023).

| IAPS Number | Content description | Mean Valence | Mean Arousal | Normative Emotion |
|---|---|---|---|---|
| 1120 | snake | 5.79 | 4.83 | fear |
| 1201 | spider | 3.47 | 5.57 | fear |
| 1300 | pitbull | 3.55 | 6.79 | fear |
| 1463 | kittens | 7.45 | 4.79 | happy |
| 1710 | puppies | 8.34 | 5.41 | happy |
| 2058 | baby | 7.91 | 5.09 | happy |
| 2165 | father | 7.63 | 4.55 | happy |
| 2205 | hospital | 1.95 | 4.53 | sad |
| 2457 | crying_boy | 3.2 | 4.94 | Sad (practice) |
| 2580 | chess | 5.71 | 2.79 | sad |
| 2800 | sad_child | 1.78 | 5.49 | sad |
| 3080 | mutilation | 1.48 | 7.22 | sad |
| 3130 | Mutilation | 1.58 | 6.97 | anger (practice) |
| 3170 | baby_tumor | 1.46 | 7.21 | sad |
| 3180 | beaten_female | 1.92 | 5.77 | anger |
| 3181 | injured_female | 2.3 | 5.06 | anger |
| 3301 | InjuredChild | 1.8 | 5.21 | sad (practice) |
| 4311 | EroticFemale | 6.66 | 6.67 | happy (practice) |
| 4533 | attractive_male | 6.22 | 5.01 | happy |
| 4599 | romance | 7.12 | 5.69 | happy |
| 4617 | flirty_female | 6.6 | 5.19 | happy |
| 4664 | erotic_female | 5.63 | 6.63 | happy |
| 4670 | erotic_couple | 6.99 | 6.74 | happy |
| 5665 | bulding | 6.15 | 4.02 | neutral |
| 5760 | nature | 8.05 | 3.22 | neutral |
| 5972 | tornado | 3.85 | 6.34 | fear |
| 6200 | aimed_gun | 2.71 | 6.21 | fear |
| 6213 | terrorist | 2.91 | 5.86 | anger |
| 6300 | knife | 2.59 | 6.61 | fear |
| 6370 | attack | 2.7 | 6.44 | fear |
| 6540 | attack | 2.19 | 6.83 | anger |
| 6571 | car_theft | 2.85 | 5.59 | fear |

| | | | | |
|---|---|---|---|---|
| 6821 | gang | 2.38 | 6.29 | anger |
| 7026 | picnic_table | 5.38 | 2.63 | neutral |
| 7354 | garlic | 5.72 | 3.5 | neutral |
| 7640 | skyscraper | 5 | 6.03 | fear |
| 8496 | waterslide | 7.58 | 5.79 | happy |
| 9181 | dead_cows | 2.26 | 5.39 | sad |
| 9253 | mutilation_girl | 2 | 5.53 | anger |
| 9265 | hung_man | 2.6 | 4.34 | sad |
| 869270 | toxic_waste | 3.72 | 5.24 | anger |
| 9400 | soldier | 2.5 | 5.99 | sad |
| 9405 | sliced_hand | 1.83 | 6.08 | fear |
| 9421 | soldier | 2.21 | 5.04 | anger |
| 9561 | sick_kitten | 2.68 | 4.79 | sad |
| 9800 | skinhead | 2.04 | 6.05 | anger |
| 9810 | kkk_rally | 2.09 | 6.62 | anger |
| 9911 | car_accident | 2.3 | 5.76 | sad |

*DIRTI Images:* Selected from the DIsgust RelaTed Images (DIRTI) database (Haberkamp et al., 2017)

| DIRTI name | Content | Mean Valence | Mean Arousal | Normative Emotion |
|---|---|---|---|---|
| 1123_body products | phlegm | 3.35 | 3.10 | disgust |
| 1131_body products | faeces | 2.74 | 3.34 | disgust |
| 1138_body products | faces | 2.07 | 4.30 | disgust |
| 1118_body products | vomit | 3.02 | 3.45 | disgust |
| 1120_body products | vomit | 2.72 | 3.62 | disgust |
| 1147_body products neutral | sink | 6.81 | 1.30 | neutral |
| 1146_body products neutral | bathroom | 6.73 | 1.37 | neutral |
| 1144_body products neutral | pavement | 5.88 | 1.23 | neutral |
| 1293_hygiene neutral | tissues | 6.65 | 1.14 | neutral |
| 1143_body products neutral | towels | 7.39 | 1.23 | neutral |

### SM2. Subjective emotion categorisation

**Fig.S1**

*Distribution of subjective emotion categories per participant*

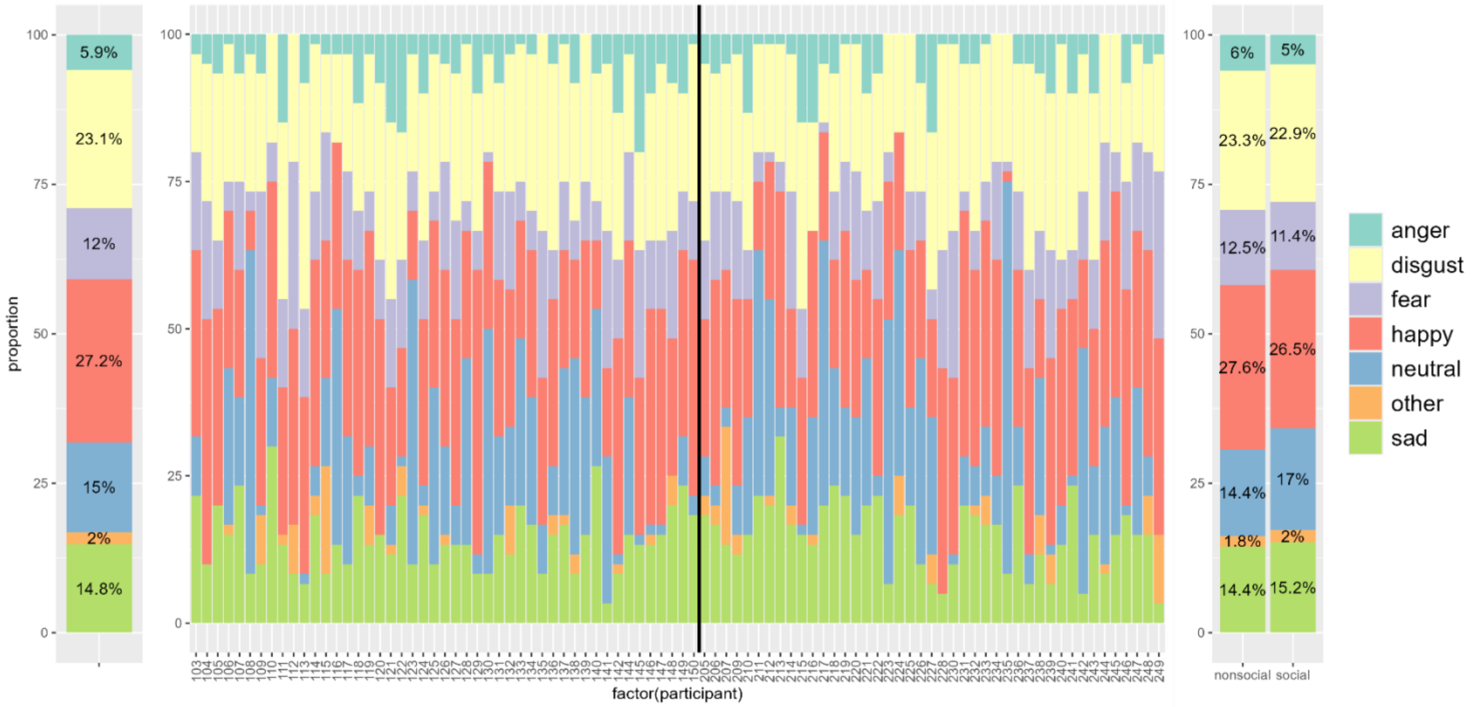

Note: (left) the total distribution of emotion categories; (middle) participant-by-participant distribution of emotion categories, with participants in the control condition displayed to the left of the black line and participants in the social observation condition displayed to the right; (right) comparison of the distribution of emotion categories across the control and social observation conditions respectively.

Given significant participant by participant variation in subjective emotion categorisation, a single emotion category was not assigned to each stimulus image (as with the normative emotion categories). Rather, the emotion category varied per participant depending on each participant's own experience of the image. That is, the emotion category used in each trial was based on the individual's experience of the stimulus not a general emotion category. The distribution of emotion categories per stimulus are displayed in Fig. S1. The subjective emotion categorisation did not always align with the normative emotion category, especially for the normative anger category. Participant reports of disgust happy were most consistent with the normative categorisation, suggesting that experience of disgust and happy are more robust than experiences of anger, fear or sad. Additionally, as there was no option for participants to explicitly report feeling neutral, neutral categorisation was inferred from low categorical emotion strength ratings, so neutral stimuli appear to have been conflated as happy. This effect was exemplified by one participant who said to the experimenter upon task completion “ordinarily a towel wouldn't make me happy, but knowing that it could have been a mutilated baby, I felt happy to see a towel!” Thus, the neutral emotion category is classed as having a positive affective valence for further analysis.

**SM3. Post-hoc analyses for results presented in section 3.1.1 – modulation of subjective affect by emotion.**

**Table S1.**

*Post-hoc comparisons of estimated marginal means for main effect of emotion on valence ratings.*

| Contrast | Estimate | SE | p |
|---|---|---|---|
| anger - disgust | -0.2201 | 0.0947 | .184 |
| anger - fear | -0.3052 | 0.0957 | .018 |
| anger - happy | -2.7355 | 0.1035 | < .001 |
| anger - neutral | -1.6025 | 0.0971 | < .001 |
| anger - sad | -0.2715 | 0.0915 | .036 |
| disgust - fear | -0.0851 | 0.0850 | .918 |
| disgust - happy | -2.5154 | 0.0890 | < .001 |
| disgust - neutral | -1.3824 | 0.0830 | < .001 |
| disgust - sad | -0.0514 | 0.0805 | .988 |
| fear - happy | -2.4304 | 0.0838 | < .001 |
| fear - neutral | -1.2973 | 0.0771 | < .001 |
| fear - sad | 0.0337 | 0.0823 | 0.999 |
| happy - neutral | 1.1330 | 0.0673 | < .001 |
| happy - sad | 2.4640 | 0.0876 | < .001 |

| | | | |
|---|---|---|---|
| neutral - sad | 1.3310 | 0.0807 | < .001 |

Note: All p values are adjusted for multiple comparisons.

**Table S2.**

*Post-hoc comparisons of estimated marginal means for main effect of emotion on intensity ratings.*

| **Contrast** | **Estimate** | **SE** | **p** |
|---|---|---|---|
| anger - disgust | 0.3825 | 0.142 | .075 |
| anger - fear | 0.4226 | 0.144 | .039 |
| anger - happy | 1.5175 | 0.154 | < .001 |
| anger - neutral | 3.0358 | 0.147 | < .001 |
| anger - sad | 0.4851 | 0.137 | .006 |
| disgust - fear | 0.0400 | 0.127 | .999 |
| disgust - happy | 1.1350 | 0.131 | < .001 |
| disgust - neutral | 2.6533 | 0.124 | < .001 |
| disgust - sad | 0.1026 | 0.120 | .957 |
| fear - happy | 1.0949 | 0.125 | < .001 |
| fear - neutral | 2.6133 | 0.117 | < .001 |
| fear - sad | 0.0625 | 0.123 | .996 |
| happy - neutral | 1.5183 | 0.102 | < .001 |
| happy - sad | -1.0324 | 0.130 | < .001 |
| neutral - sad | -2.5507 | 0.121 | < .001 |

Note: All p values are adjusted for multiple comparisons.

**Table S3.** Post-hoc comparisons of estimated marginal means for interaction effect of emotion and social context on intensity ratings:

| **Emotion** | **Contrast (Non-social − Social)** | **Estimate** | **SE** | **p** |
|---|---|---|---|---|
| anger | non-social − social | 0.1865 | 0.292 | .523 |
| disgust | non-social − social | 0.1913 | 0.227 | .399 |
| fear | non-social − social | 0.3394 | 0.251 | .177 |
| happy | non-social − social | 0.6065 | 0.224 | .007 |
| neutral | non-social − social | 0.2548 | 0.245 | .299 |
| sad | non-social − social | -0.0644 | 0.240 | .789 |

Note: All p values are adjusted for multiple comparisons.

### SM4. Post-hoc analyses for results presented in section 3.1.3 – modulation of facial response components by emotion and social context

**Table S4.**

*Post-hoc comparisons of estimated marginal means for main effect of emotion on C1.*

| Contrast | Estimate | SE | p |
|---|---|---|---|
| anger - disgust | -0.51693 | 0.176 | .039 |
| anger - fear | -0.05484 | 0.190 | .999 |
| anger - happy | -0.18183 | 0.177 | .909 |
| anger - neutral | -0.06428 | 0.189 | .999 |
| anger - sad | -0.02572 | 0.182 | 1.000 |
| disgust - fear | 0.46209 | 0.142 | .015 |
| disgust - happy | 0.33511 | 0.119 | .055 |
| disgust - neutral | 0.45265 | 0.136 | .012 |
| disgust - sad | 0.49121 | 0.133 | .003 |
| fear - happy | -0.12698 | 0.139 | .944 |
| fear - neutral | -0.00944 | 0.154 | 1.000 |
| fear - sad | 0.02912 | 0.152 | 1.000 |
| happy - neutral | 0.11754 | 0.130 | .945 |
| happy - sad | 0.15611 | 0.131 | .843 |
| neutral - sad | 0.03856 | 0.146 | .999 |

Note: All p values are adjusted for multiple comparisons.

**Table S5.**

*Post-hoc comparisons of estimated marginal means for main effect of emotion on C2.*

| Contrast | Estimate | SE | p |
|---|---|---|---|
| anger - disgust | -0.28247 | 0.158 | .470 |
| anger - fear | 0.04304 | 0.168 | .999 |
| anger - happy | 0.19231 | 0.160 | .836 |
| anger - neutral | 0.21250 | 0.167 | .802 |
| anger - sad | 0.03818 | 0.161 | .999 |
| disgust - fear | 0.32551 | 0.130 | .121 |
| disgust - happy | 0.47478 | 0.113 | 0.000 |

| | | | |
|---|---|---|---|
| disgust - neutral | 0.49498 | 0.124 | 0.001 |
| disgust - sad | 0.32066 | 0.122 | 0.090 |
| fear - happy | 0.14927 | 0.127 | 0.849 |
| fear - neutral | 0.16947 | 0.136 | 0.814 |
| fear - sad | -0.00485 | 0.136 | 1.000 |
| happy - neutral | 0.02020 | 0.115 | 1.000 |
| happy - sad | -0.15412 | 0.122 | .806 |
| neutral - sad | -0.17432 | 0.131 | .769 |

Note: All p values are adjusted for multiple comparisons.

**Table S6**.

*Post-hoc comparisons of estimated marginal means for main effect of emotion on C3.*

| Contrast | Estimate | SE | p |
|---|---|---|---|
| anger - disgust | 0.2257 | 0.171 | .772 |
| anger - fear | 0.0627 | 0.184 | .999 |
| anger - happy | -0.1081 | 0.171 | .989 |
| anger - neutral | -0.1845 | 0.183 | .915 |
| anger - sad | -0.0482 | 0.177 | .999 |
| disgust - fear | -0.1630 | 0.137 | .844 |
| disgust - happy | -0.3338 | 0.115 | .044 |
| disgust - neutral | -0.4102 | 0.132 | .023 |
| disgust - sad | -0.2739 | 0.128 | .270 |
| fear - happy | -0.1708 | 0.135 | .803 |
| fear - neutral | -0.2472 | 0.149 | .558 |
| fear - sad | -0.1109 | 0.147 | .975 |
| happy - neutral | -0.0764 | 0.125 | .990 |
| happy - sad | 0.0599 | 0.127 | .997 |
| neutral - sad | 0.1363 | 0.141 | .929 |

Note: All p values are adjusted for multiple comparisons.

**Table S7.**

*Post-hoc comparisons of estimated marginal means for interaction effect of emotion and social context on C2.*

| Emotion | Contrast (Non-social − Social) | Estimate | SE | p |
|---|---|---|---|---|
| anger | non-social − social | 1.039 | 0.579 | .072 |
| disgust | non-social − social | 0.575 | 0.533 | .280 |
| fear | non-social − social | 0.537 | 0.559 | .328 |
| happy | non-social − social | 1.258 | 0.531 | .017 |
| neutral | non-social − social | 1.149 | 0.545 | .035 |
| sad | non-social − social | 1.169 | 0.541 | .031 |

Note: All p values are adjusted for multiple comparisons.

**SM5. Dimensional analysis of subjective, autonomic and behavioural affective response in long format (Observations > features)**

The figure below summarises the results of the MFA cross-modal dimension analysis where dimension one capturing higher contributions from variables across modalities on average is taken as an index of coherence. This was conducted by treating both participant and trials observations (rows), and autonomic, subjective and behavioural measures as variable groups (columns). Note that while this was pre-registered, and increases power compared to the main analysis reported in the paper, it nonetheless risks inflating differences in variables of interest due to mixing within and between subject variances (see Methods in the manuscript).

**Fig S2.**

*MFA Sensitivity analysis in long (repeated measures) format*

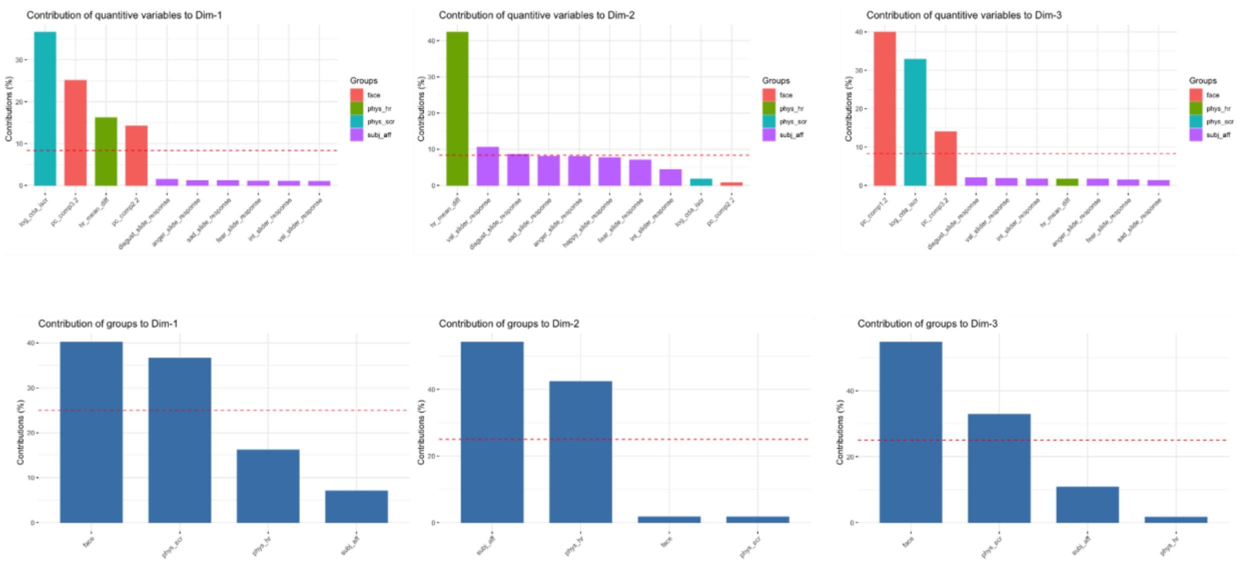

Note: Phys_hr – Heart rate change; Phys_scr = Skin Conductance Response; Subj_aff = subjective affective responses (valence and emotion ratings), face = facial behaviour components.

Tables S8-S10 summarises the mixed-effects model analyses on the covariance-derived dimensions for subjective, autonomic, and behavioural affective responses. Dimension 1 reflects contributions from most affective modalities (Fig S2) and is therefore interpreted as indexing coherence consistent with the main results in the manuscript (3.2). The effect of social context on this dimension is consistent with the main results reported in the paper, showing differences in cross-modal coherence between the social compared to the non-social condition (Table S8).

For simplicity of presentation, the tables report estimates using neutral emotion as the reference category for the subjective emotion factor. Notably, the results also show strong effects of emotion on coherence. Effects observed on the remaining dimensions are largely driven by emotion consistent with unimodal patterns, which were themselves primarily influenced by subjective emotion (3.1)

**Table S8**.

*Summary of mixed model analysis on dimension 1.*

| | **mfa pc 1** | | |
|---|---|---|---|
| *Predictors* | *Estimates* | *CI* | *p* |
| (Intercept) | 0.09 | -0.13 – 0.31 | .416 |
| social nonsocial [social] | -0.55 | -0.86 – -0.24 | **<.001** |
| subjective emotion [anger] | 0.38 | 0.24 – 0.52 | **<.001** |
| subjective emotion [disgust] | 0.33 | 0.22 – 0.43 | **<.001** |
| subjective emotion [fear] | 0.25 | 0.14 – 0.37 | **<.001** |
| subjective emotion [happy] | -0.14 | -0.24 – -0.04 | **.006** |
| subjective emotion [sad] | 0.24 | 0.13 – 0.35 | **<.001** |
| social nonsocial [social] × subjective emotion [anger] | 0.04 | -0.16 – 0.23 | .716 |
| social nonsocial [social] × subjective emotion [disgust] | 0.13 | -0.01 – 0.26 | .063 |
| social nonsocial [social] × subjective emotion [fear] | 0.11 | -0.05 – 0.27 | .177 |
| social nonsocial [social] × subjective emotion [happy] | 0.09 | -0.04 – 0.22 | .190 |
| social nonsocial [social] × subjective emotion [sad] | 0.13 | -0.02 – 0.27 | .098 |
| **Random Effects** | | | |
| $\sigma^2$ | 0.54 | | |
| $\tau_{00\ participant}$ | 0.49 | | |
| $\tau_{00\ stim\_iaps}$ | 0.01 | | |
| ICC | 0.48 | | |
| $N_{participant}$ | 90 | | |
| $N_{stim\_iaps}$ | 60 | | |
| Observations | 5400 | | |
| Marginal $R^2$ / Conditional $R^2$ | 0.087 / 0.529 | | |

**Table S9**.

*Summary of mixed model analysis on dimension 2.*

| | **mfa pc 2** | | |
|---|---|---|---|
| *Predictors* | *Estimates* | *CI* | *p* |
| (Intercept) | -0.37 | -0.52 – -0.22 | **<.001** |
| social nonsocial [social] | 0.10 | -0.06 – 0.26 | .207 |
| subjective emotion [anger] | 1.02 | 0.89 – 1.15 | **<.001** |
| subjective emotion [disgust] | 0.67 | 0.57 – 0.78 | **<.001** |
| subjective emotion [fear] | 0.62 | 0.51 – 0.72 | **<.001** |
| subjective emotion [happy] | -0.17 | -0.26 – -0.07 | **<.001** |
| subjective emotion [sad] | 0.62 | 0.51 – 0.73 | **<.001** |
| social nonsocial [social] × subjective emotion [anger] | -0.10 | -0.28 – 0.07 | .253 |
| social nonsocial [social] × subjective emotion [disgust] | -0.06 | -0.18 – 0.06 | .347 |
| social nonsocial [social] × subjective emotion [fear] | -0.06 | -0.21 – 0.08 | .399 |
| social nonsocial [social] × subjective emotion [happy] | -0.06 | -0.18 – 0.07 | .367 |
| social nonsocial [social] × subjective emotion [sad] | -0.06 | -0.20 – 0.07 | .347 |
| **Random Effects** | | | |
| $\sigma^2$ | 0.45 | | |
| $\tau_{00\ participant}$ | 0.09 | | |
| $\tau_{00\ stim\_iaps}$ | 0.14 | | |
| ICC | 0.34 | | |
| $N_{participant}$ | 90 | | |
| $N_{stim\_iaps}$ | 60 | | |
| Observations | 5400 | | |
| Marginal $R^2$ / Conditional $R^2$ | 0.192 / 0.467 | | |

**Table S10.**

*Summary of mixed model analysis on dimension 3.*

| | **mfa pc 3** | | |
|---|---|---|---|
| *Predictors* | *Estimates* | *CI* | *p* |
| (Intercept) | -0.23 | -0.45 – -0.01 | **.043** |
| social nonsocial [social] | 0.10 | -0.21 – 0.42 | .521 |
| subjective emotion [anger] | 0.43 | 0.32 – 0.54 | **<.001** |
| subjective emotion [disgust] | 0.44 | 0.35 – 0.53 | **<.001** |
| subjective emotion [fear] | 0.34 | 0.25 – 0.44 | **<.001** |
| subjective emotion [happy] | -0.13 | -0.22 – -0.05 | **.001** |
| subjective emotion [sad] | 0.31 | 0.22 – 0.41 | **<.001** |
| social nonsocial [social] × subjective emotion [anger] | 0.01 | -0.15 – 0.17 | 0.881 |
| social nonsocial [social] × subjective emotion [disgust] | -0.01 | -0.12 – 0.10 | .873 |
| social nonsocial [social] × subjective emotion [fear] | -0.07 | -0.21 – 0.06 | .280 |
| social nonsocial [social] × subjective emotion [happy] | -0.02 | -0.13 – 0.09 | .700 |
| social nonsocial [social] × subjective emotion [sad] | 0.02 | -0.10 – 0.14 | .754 |
| **Random Effects** | | | |
| $\sigma^2$ | 0.37 | | |
| $\tau_{00\ \text{participant}}$ | 0.53 | | |
| $\tau_{00\ \text{stim\_iaps}}$ | 0.01 | | |
| ICC | 0.60 | | |
| $N_{\text{participant}}$ | 90 | | |
| $N_{\text{stim\_iaps}}$ | 60 | | |
| Observations | 5400 | | |
| Marginal $R^2$ / Conditional $R^2$ | 0.062 / 0.621 | | |

**SM6. Subject-level multiple factor analysis.**

As an additional sensitivity analysis for the coherence results, we derived MFA solutions at the participant level. This approach was motivated by the possibility that the main analyses may be underpowered, given the relatively large number of variables (trials and measures treated as variables) in relation to observations (participants) in the main MFA reported in the paper. Additionally, group-level analysis may not fully capture individual-specific covariance patterns. In particular, if individuals differ systematically in their covariance structure for autonomic, subjective and behavioural affective responses across trials, the group-level MFA, by imposing a shared dimensional structure may obscure differences between social and non-social conditions). While multilevel extensions of dimensionality reduction techniques have been proposed (Abdi et al., 2013; Husson et al., 2017; Rohart et al., 2017), these methods are not straightforward to apply in the context of MFA, and current implementations typically rely on hierarchical groupings of variables rather than by participants or vice-versa.

To address this limitation, we implemented a custom subject-level MFA procedure in which separate MFA models were estimated for each participant, allowing the dimensional structure to vary across individuals. This approach enables a more direct characterisation of individual-specific covariance patterns and provides a complementary perspective on the coherence results reported in the main analyses. The pipeline involved the following steps:

1. Group-level MFA. A multiple factor analysis (MFA) was performed on the full dataset to derive a group-level latent structure across all modalities (e.g.

subjective ratings, heart rate, skin conductance and facial action units). This was done to provide an overall view of the covariance for latter steps

**Fig.S3**

*Modality contributions to group-level MFA dimensions*

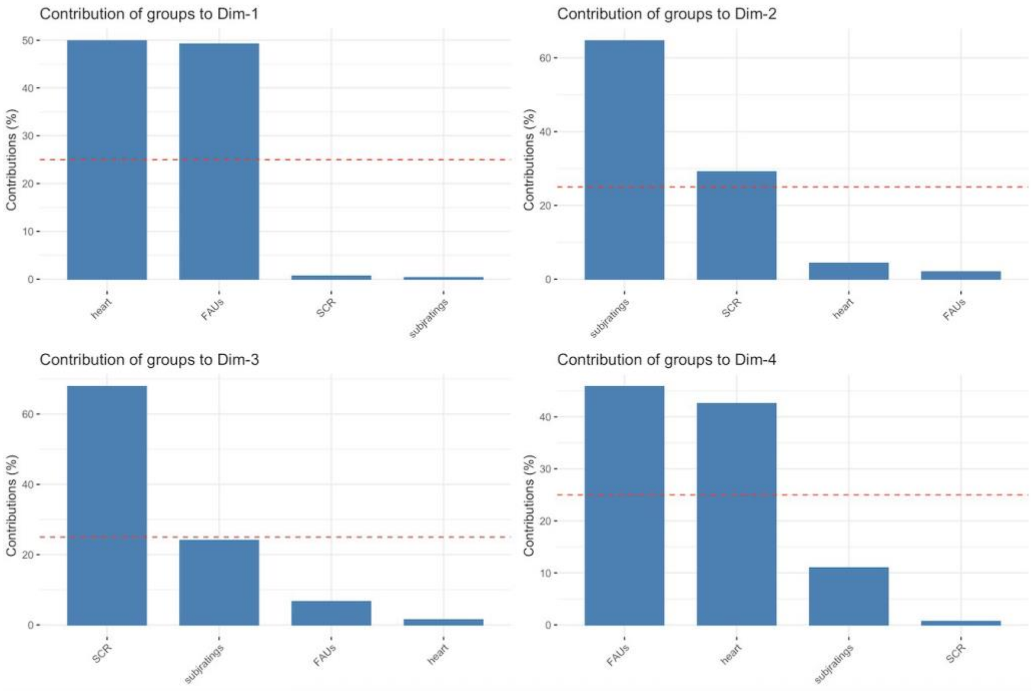

Note: Heart = Heart rate change; SCR = Skin Conductance Response; subj ratings = subjective affective responses (valence and emotion ratings), FAUs = facial action units.

**Fig.S4**

*Individual variable contributions to group-level MFA dimensions*

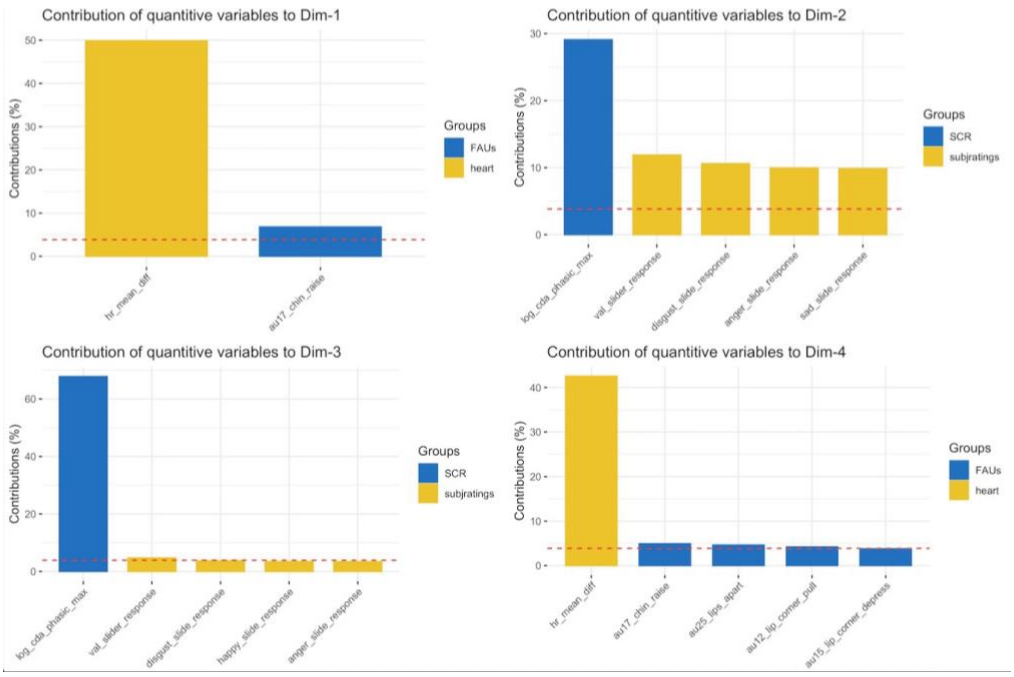

Note: Heart = Heart rate change; SCR = Skin Conductance Response; subj ratings = subjective affective responses (valence and emotion ratings), FAUs= facial action units.

Note: Heart = Heart rate change; SCR = Skin Conductance Response; subj ratings = subjective affective responses (valence and emotion ratings), FAUs= facial action units.

2. Participant-level MFAs. Separate MFAs were conducted for each participant using the same variable grouping structure. This gave participant-specific latent dimensions, variable loadings and variance explained for each group of variables (e.g. subjective ratings) and individual variables (e.g., valence rating).

3. Alignment to group structure. Because the subject specific MFAs might have very different structures, to aggregate results across groups and compare groups, we needed to align the subject specific. This was done by aligning it to the group solution (in 1) using Procrustes rotation (using the R package “vegan” (Oksanen et al., 2019) which minimises discrepancies between individual and group configurations through rotation of one set of data to a target data set.
4. Validation of alignment. Correlations between participant-averaged MFAs and the group MFA were computed before and after Procrustes transformation, confirming improved correspondence following alignment.

**Fig.S5**

*Pre- and post-alignment correlation matrices*

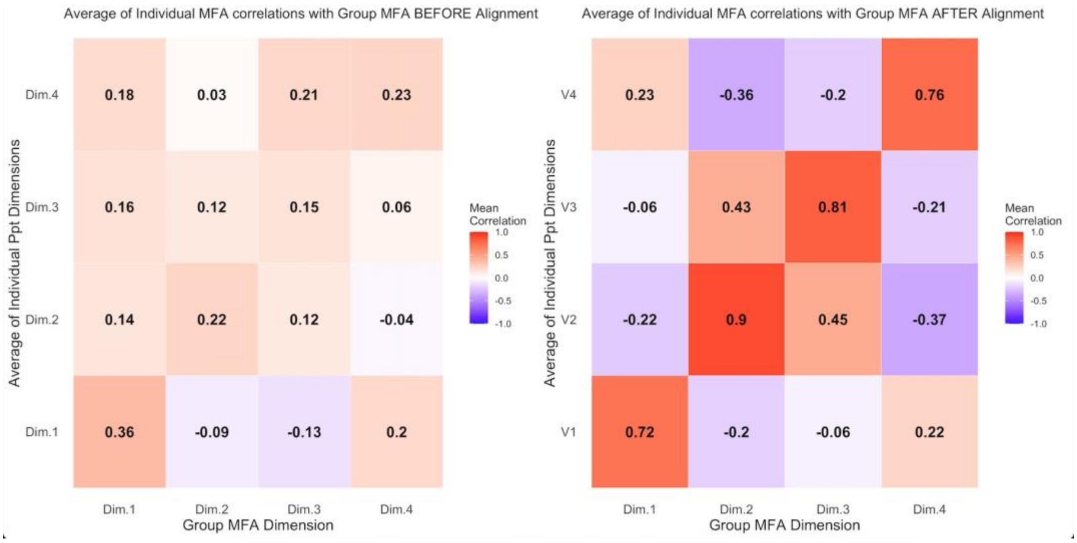

5. Coherence metrics. Following alignment, metrics of structural coherence were derived, including the variance explained by the first aligned dimension and the cumulative variance explained by the first four dimensions. These metrics were subsequently compared across social and non-social contexts.

**Table S11.**

*Metric comparisons between social and non-social contexts*

| Measure | Non-social | Social | *t* | df | *p* |
|---|---|---|---|---|---|
| Dim 1 Variance (%) | 19.50 | 17.32 | 1.74 | 88.0 | .085 |
| Var explained by Dims 1-4 | 60.09 | 59.90 | 0.12 | 86.5 | .906 |

Note: Degrees of freedom are approximated via Welch correction for inequality of variances.

Results indicate that there isn't a fundamental difference in structural coherence (i.e. covariance structure) between social vs non-social groups, despite the observed difference along the coherence dimensions reported in 3.2, and Table S8.

**SM7. Mixed models analysis of gaze metrics.**

**Table S12.**

*Summary of mixed model analysis for fixation duration, saccade amplitudes and entropy.*

| Measure | Effect | *df* (Num, Den) | *F* | *p* |
|---|---|---|---|---|
| **Fixation duration** | Social nonsocial | 1, 97.7 | 0.90 | .346 |
| | Emotion | 5, 1546.1 | 1.12 | .346 |
| | Social × Emotion | 5, 5149.2 | 0.51 | .769 |
| **Saccade amplitude** | Social nonsocial | 1, 95.3 | 0.00 | .955 |
| | Emotion | 5, 4981.1 | 0.12 | .989 |
| | Social × Emotion | 5, 5124.6 | 1.56 | .167 |
| **Entropy** | Social nonsocial | 1, 96.2 | 1.91 | .170 |
| | Emotion | 5, 4793.2 | 1.40 | .219 |
| | Social × Emotion | 5, 5130.3 | 1.93 | .087 |

Note: Type III analysis of variance for fixed effects from linear mixed-effects models (Satterthwaite approximation for degrees of freedom).

**SM8. Model comparisons**

In response to reviewer feedback, we compare here linear mixed effects models that included emotion as random slopes with our original models reported in the manuscript. The models had the following format:

Outcome ~ SocialContext * SubjectiveEmotionCategory + (1 + SubjectiveEmotionCategory | Participant) + (1 | Stimulus)

The requested random-slope refits produced convergence warnings, singular fits, and/or evidence that the additional complexity was not reliably supported by the data (based on information criteria, AIC, BVI, LRT). These issues were not resolved by increased optimiser iterations, alternative optimisers, checking maximum Hessian gradients and fitting uncorrelated random-effects structures. We therefore do not treat these models as a stable basis for replacing or reinterpreting the primary analyses. Importantly, the requested refits did not suggest a reversal of the effects reported in the manuscript.

**Table S13.**

*Comparison of reported and random-slope refit models*

| **Outcome** | **Random-slope refit** | **Diagnostics** | **Model comparison** | **Conclusion** |
|---|---|---|---|---|
| **Valence** | Emotion effect retained; interaction effect remained n.s; post-hoc pattern unchanged. | Convergence warning; Hessian gradient issue. | ΔAIC = -838, ΔBIC = -706; LRT p <.001 | Improved fit but unresolved convergence issues; no change in inference. |

| | | | | |
|---|---|---|---|---|
| **Intensity** | Emotion effect retained; social context n.s; omnibus interaction became non-significant, but the reported interaction post-hoc retained the same direction and significance. | Singular fit. | ΔAIC = -160; ΔBIC = -28; LRT p <.001 | Improved fit but singularity indicates unstable random slope structure. |
| **Heart rate** | Social context effect retained; emotion and interaction n.s; emotion contrasts remained n.s | Singular fit. | ΔAIC = +27; ΔBIC = +160; LRT p =.881 | Added complexity not supported. |
| **SCR** | No significant social context, emotion, or interaction effects. | Singular fit. | ΔAIC = +30; ΔBIC = +161; LRT p =.951 | Added complexity not supported. |
| **Face C2** | Emotion and interaction effects retained. | Singular fit. | ΔAIC = +10; ΔBIC = +141; LRT p =.070 | Added complexity not supported. |

Note: n.s. = non significant. **Note**. n.s. = non-significant. ΔAIC, Akaike Information Criteria change, ΔBIC; Bayesian Information Criteria change, LRT: Likelihood Ratio Test

We also conducted supplementary analyses using AU-emotion groupings suggested by our reviewer: AU6 + AU12 for happiness, AU4 + AU5 + AU7 + AU23 for anger, AU1 + AU4 + AU15 for sadness, AU1 + AU2 + AU5 + AU26 for fear, and AU9 + AU10 for disgust.  For example, the model for the happiness-related AU grouping was:

AU6_AU12 activity ~ social context * subjective emotion category + (1 | participant) + (1 | stimulus)

Consistent with the original component-based analyses, several AU-emotion groupings showed significant main effects of subjective emotion category. This indicates that the selected AU groupings captured some emotion-related variation in facial activity. However, these effects did not map exclusively onto the intended emotion category. For example, as in the component-based results, disgust was among the more consistently differentiated subjective emotion categories across the AU-group analyses (which is consistent with our reported component results). Importantly, none of the AU-grouping models showed a significant subjective emotion × social context interaction. Thus, the requested AU-group analyses partly recovered emotion-related differences in facial activity, but they did not capture the social modulation observed in the component-based facial analysis.

**Table S14**

*Facial analysis by emotion specific action units.*

| Outcome | Predictors | F(df1, df2) | *p* |
|---|---|---|---|
| Happiness (AU12, AU6) | Social context | F(1, 92.2) = .041 | .84 |
| | Emotion | F(5, 524.2) = 2.71 | **.02** |
| | Social x Emotion | F(5, 5366.9) = 1.17 | .32 |
| Anger (AU04, AU05, AU07, AU23) | Social context | F(1, 92.6) = 1.85 | .18 |
| | Emotion | F(5, 736.3) = 3.29 | **.01** |
| | Social x Emotion | F(5, 5369.2) = .29 | .91 |
| Sadness (AU01, AU04, AU15) | Social context | F(1, 93.1) = .32 | .58 |
| | Emotion | F(5, 748.5) = 5.07 | **<.001** |
| | Social x Emotion | F(5, 5371.4) = 1.04 | .39 |
| Fear (AU01, AU02, AU05, AU26) | Social context | F(1, 94.2) = .00 | .94 |
| | Emotion | F(5, 530.8) = 1.07 | .38 |
| | Social x Emotion | F(5, 5393.9) = .31 | .91 |
| Disgust (AU09, AU10) | Social context | F(1, 92.7) = .73 | .39 |
| | Emotion | F(5, 478) = 4.21 | **<.001** |
| | Social x Emotion | F(5, 5376.7) = 1.15 | .33 |

Note: Type III analysis of variance for fixed effects from linear mixed-effects models (Satterthwaite approximation for degrees of freedom).

**Table S15**

*Post-hoc comparisons of estimated marginal means for interaction effect of emotion and social context on C2.*

| Outcome | Contrast | Estimate | p |
|---|---|---|---|
| Happiness (AU12, AU6) | anger - disgust | -0.05857 | .313 |
| | anger - fear | -0.00336 | 1 |
| | anger - happy | -0.01217 | .998 |
| | anger - neutral | 0.01032 | .999 |
| | anger - sad | 0.00224 | 1 |
| | disgust - fear | 0.05520 | .161 |
| | disgust - happy | 0.04640 | .167 |
| | disgust - neutral | 0.06889 | **.024** |
| | disgust - sad | 0.06081 | .057 |
| | fear - happy | -0.00881 | .999 |
| | fear - neutral | 0.01368 | .994 |
| | fear - sad | 0.00560 | .999 |
| | happy - neutral | 0.02249 | .893 |
| | happy - sad | 0.01441 | .985 |
| | neutral - sad | -0.00808 | .999 |
| Anger (AU04, AU05, AU07, AU23) | anger - disgust | -0.01642 | .988 |
| | anger - fear | -0.01339 | .997 |
| | anger - happy | 0.03971 | .652 |
| | anger - neutral | 0.04796 | .497 |
| | anger - sad | 0.01337 | .996 |
| | disgust - fear | 0.00302 | 1 |
| | disgust - happy | 0.05613 | **.028** |
| | disgust - neutral | 0.06438 | **.019** |
| | disgust - sad | 0.02978 | .670 |
| | fear - happy | 0.05311 | .108 |
| | fear - neutral | 0.06135 | .065 |
| | fear - sad | 0.02676 | .837 |
| | happy - neutral | 0.00825 | .998 |
| | happy - sad | -0.02635 | .775 |
| | neutral - sad | -0.03459 | .592 |
| Sadness | anger - disgust | -0.08427 | .100 |

| | | | |
|---|---|---|---|
| (AU01, AU04, AU15) | anger - fear | 0.02480 | .980 |
| | anger - happy | 0.01586 | .997 |
| | anger - neutral | 0.01112 | .999 |
| | anger - sad | -0.00567 | 1 |
| | disgust - fear | 0.10907 | **.001** |
| | disgust - happy | 0.10013 | **.000** |
| | disgust - neutral | 0.09539 | **.003** |
| | disgust - sad | 0.07859 | **.024** |
| | fear - happy | -0.00894 | .999 |
| | fear - neutral | -0.01369 | .997 |
| | fear - sad | -0.03048 | .889 |
| | happy - neutral | -0.00474 | 1 |
| | happy - sad | -0.02153 | .959 |
| | neutral - sad | -0.01679 | .989 |
| Fear (AU01, AU02, AU05, AU26) | anger - disgust | -0.05158 | .767 |
| | anger - fear | -0.02644 | .988 |
| | anger - happy | 0.00055 | 1 |
| | anger - neutral | -0.00223 | 1 |
| | anger - sad | -0.00400 | 1 |
| | disgust - fear | 0.02515 | .967 |
| | disgust - happy | 0.05213 | .356 |
| | disgust - neutral | 0.04935 | .569 |
| | disgust - sad | 0.04758 | .585 |
| | fear - happy | 0.02699 | .951 |
| | fear - neutral | 0.02421 | .980 |
| | fear - sad | 0.02244 | .985 |
| | happy - neutral | -0.00278 | 1 |
| | happy - sad | -0.00455 | 1 |
| | neutral - sad | -0.00177 | 1 |
| Disgust (AU09, AU10) | anger - disgust | -0.09553 | .0734 |
| | anger - fear | -0.02703 | .9801 |
| | anger - happy | 0.00240 | 1 |
| | anger - neutral | -0.00700 | 1 |
| | anger - sad | -0.01297 | .999 |
| | disgust - fear | 0.06850 | .154 |
| | disgust - happy | 0.09793 | **.001** |
| | disgust - neutral | 0.08853 | **.015** |
| | disgust - sad | 0.08256 | **.023** |
| | fear - happy | 0.02943 | .899 |
| | fear - neutral | 0.02003 | .987 |
| | fear - sad | 0.01406 | .997 |
| | happy - neutral | -0.00939 | .999 |
| | happy - sad | -0.01537 | .992 |
| | neutral - sad | -0.00597 | 1 |

Note: All p values are adjusted for multiple comparisons.

Finally, we also fitted restricted models in which each AU grouping (e.g. AU6 + AU12 for happy) was analysed only within the corresponding subjective emotion category. In other words, AU6 + AU12 would be analysed only for trials subjectively categorised as happy, AU4 + AU5 + AU7 + AU23 only for trials subjectively categorised as anger, and so forth. We therefore also fitted these models as an additional sensitivity analysis. In these models, subjective emotion category is necessarily removed as a predictor because each model includes only one emotion category.

For example, the model for the happiness-related AU grouping was:

AU6_AU12 activity ~ social context + (1 | participant) + (1 | stimulus)

The models therefore tested whether social context modulated the target AU grouping within the theoretically corresponding emotion category. The results of these restricted models were consistent with the broader AU-grouping analyses. There was no reliable evidence that social context modulated the predefined AU groupings within their corresponding subjective emotion categories.

**Table S16**

| **Outcome** | **Predictor** | **F(df1, df2)** | ***p*** |
|---|---|---|---|
| Happiness<br>(AU12, AU6) | Social context | F(1, 92.1) = 006 | .80 |
| Anger<br>(AU04, AU05, AU07, AU23) | Social context | F(1, 74.4) = 2.96 | .08 |
| Sadness<br>(AU01, AU04, AU15) | Social context | F(1, 89.6) = .15 | .69 |
| Fear<br>(AU01, AU02, AU05, AU26) | Social context | F(1, 94.0) = .00 | .94 |
| Disgust | Social context | F(1, 91.4) = .27 | .60 |

| |
|---|
| (AU09, AU10) |

Note: Type III analysis of variance for fixed effects from linear mixed-effects models (Satterthwaite approximation for degrees of freedom).

**SM9. Reduced-feature MFA sensitivity analysis of cross-modal affective coherence**

As an additional reduced-feature sensitivity analysis, we repeated the wide-format MFA after aggregating responses across the experimentally selected emotion categories while retaining the same 12 response features (across subjective affect and emotion categories, physiology and face components). This analysis was intended to address concerns about the relatively large number of features in the primary MFA as well the non-independence of a long-format MFA. Aggregating across emotion categories reduced the input matrix from 90 participants × 720 trial-level response variables to 90 participants × 72 emotion-averaged response variables. We first inspected the group contributions to Dimension 1 to confirm that it retained a cross-modal structure comparable to the primary MFA solution. Dimension 1 was therefore treated as the reduced-feature index of cross-modal affective coherence.

The reduced-feature MFA reproduced the main coherence result: participants in the non-social condition scored significantly higher on Dimension 1 than participants in the social condition, $t(85.67) = 3.74$, $p < .001$, $d = 0.79$, with higher mean scores in the non-social group ($M = 0.42$) than in the social group ($M = -0.46$). In contrast, participant contributions to Dimension 1 did not differ significantly between conditions, $t(82.50) = -0.52$, $p = .607$, indicating that the group difference was not attributable to unequal representation of the latent dimension across conditions. The consistency between the original wide-format MFA (with trial and variables as features), the long-format MFA (with participant and trial level as observations), and the present reduced-feature MFA therefore supports the robustness of the main coherence finding and suggests that it was not driven by instability arising from the large trial-level feature space.

**SM10. Simulation-based power analysis for unimodal analyses**

Our primary hypothesis concerned the between-participant social versus non-social contrast. We therefore treated this contrast as the key limiting factor for power and evaluated it using a simulation-based sensitivity analysis that matched the core structure of the reported unimodal models. Prior work has reported small to moderate social context effects on facial and gestural emotional signalling (Heesen et al., 2024) including minimal effects of "being watched" on facial EMG and autonomic responding (Hietanen et al., 2019). At the same time, experimental and meta-analytic evidence suggests that autonomic effects are heterogeneous and often modest rather than large and uniform across contexts (Cuve et al., 2023; Siegel et al., 2018). Given the subtle and non-interactive nature of the present manipulation, we therefore treated a small but meaningful effect as the representative unimodal social-context effect of interest. The assumed effect was chosen to be broadly comparable and approximate a small Cohen's d (Cohen, 1988; Lakens, 2013), specified a fixed effect of 0.30 on a standardised outcome scale.

However, an exact a-priori power analysis for the full crossed random-effects model was not feasible without specifying several parameters that are not known with confidence in advance, including participant- and stimulus-level variance components, residual variance, and the correlation structure among repeated observations as well as the fact that some fixed effects are data-driven (e.g. subjectively derived emotion categories, which constitute an additional source of variance as discussed earlier). We therefore used an approximation based on simulation analysis conducted in simr (Green & MacLeod, 2016; Kumle et al., 2021). Specifically, we conducted 100 simulations of data under a model of the form:

$$y \sim \text{social} * \text{emotion} + (1 \mid \text{participant}) + (1 \mid \text{stim})$$

The simulated design approximated the unimodal structure and matched the study in broad terms (N = 90; 45 per group, 60 stimuli; 6 emotion categories). Under the assumed parameter values, estimated power for the primary social-context contrast was 95.0% (95% CI [88.72, 98.36]). Furthermore, the corresponding power curve indicated that power was below 80% at N = 40 but above 80% by N = 60 (see power curve below). Accordingly, we interpret these analyses primarily as evidence that the study was adequately powered to the broad aims of exploring a social modulation of affective responses.

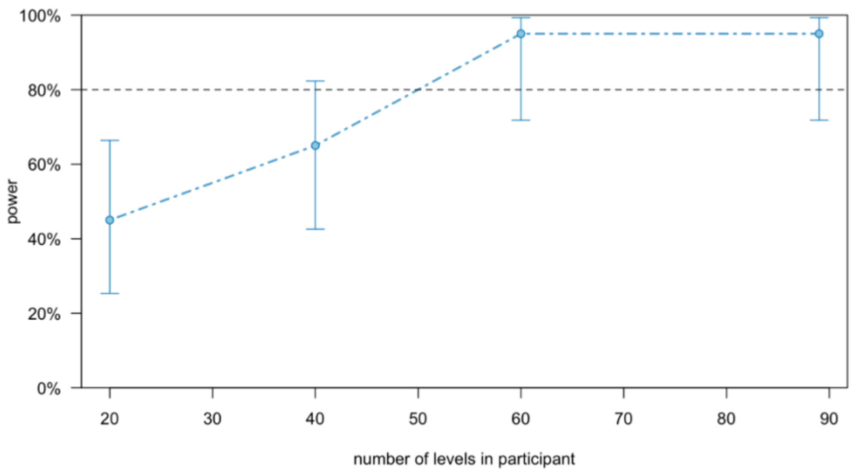

At the same time, we note that precise a-priori power estimation for all possible contrasts and interaction terms in the full mixed-effects model is not practical without making substantial assumptions.
